# Carbon Nanocarriers Deliver siRNA to Intact Plant Cells for Efficient Gene Knockdown

**DOI:** 10.1101/564427

**Authors:** Gozde S. Demirer, Huan Zhang, Natalie S. Goh, Rebecca L. Pinals, Roger Chang, Markita P. Landry

## Abstract

Post-transcriptional gene silencing (PTGS) is a powerful tool to understand and control plant metabolic pathways, which is central to plant biotechnology. PTGS is commonly accomplished through delivery of small interfering RNA (siRNA) into cells. Standard plant siRNA delivery methods (*Agrobacterium* and viruses) involve coding siRNA into DNA vectors, and are only tractable for certain plant species. Herein, we develop a nanotube-based platform for direct delivery of siRNA, and show high silencing efficiency in intact plant cells. We demonstrate that nanotubes successfully deliver siRNA and silence endogenous genes owing to effective intracellular delivery and nanotube-induced protection of siRNA from nuclease degradation. This study establishes that nanotubes could enable a myriad of plant biotechnology applications that rely on RNA delivery to intact cells.

## INTRODUCTION

Plants are central in providing over 25% of our most clinically-relevant drugs, are at the core of our sustainability efforts, and will benefit from genetic engineering to feed our growing population in the midst of climate change. Plant biotechnology is currently limited by the cost, ease, and throughput of methods for probing plant genetics, and by the complexity of plant biosynthetic pathways. Consequently, less than a dozen complete biosynthetic pathways are known for plant natural products that have been reconstituted heterologously, compared to the ~1000 known biosynthetic pathways in bacteria and fungi (*1*). RNA interference (RNAi) is sequence-specific inhibition of gene expression at the messenger RNA (mRNA) level(*2*), and can either consist of transcriptional gene silencing (TGS) or post-transcriptional gene silencing (PTGS). In PTGS, small RNA molecules – micro (miRNA) or small interfering (siRNA) – direct enzyme complexes to degrade mRNA molecules, hence suppress their activity by preventing translation.

PTGS has shown to be a prominent tool in plants for genotype-phenotype mapping (*3*), discovery of new biosynthetic pathways (*4, 5*), increased production of valuable small molecules (*6, 7*), understanding the functions of genes and proteins (*8*), and to confer resistance to plant diseases (*9–11*). One common way of utilizing PTGS in plants is to directly deliver siRNA molecules into cells. However, plants have a cell wall which presents a barrier to exogenous biomolecule delivery, whereby the plant cell wall size exclusion limit is ~ 5-20 nm (*12*). Consequently, viral vectors combined with *Agrobacterium tumefaciens* delivery is the preferred method to deliver siRNA into intact plant cells. Viral vectors present the advantage of directly and strongly expressing the siRNA without relying on plant transformation, however, most viruses are limited in their host range (*13*), often do not result in uniform silencing of the gene, and thus levels of silencing can vary between plants and experiments (*14*), and might inadvertently result in the suppression of non-target genes. *Agrobacterium-*mediated delivery, similarly, is also limited to use in certain plant species, often yields random DNA integration that can adversely and unpredictably affect the cell operation (*15*), results in constitutive expression of siRNA thus limiting temporal control over gene silencing, and can be difficult to scale or multiplex for high-throughput or multi-gene target applications, respectively (*16*).

While nanomaterial-mediated delivery of RNA and therapeutics has been extensively explored in animals (*17–19*), its potential for plant systems remains under-studied (*20*). Several prior studies take advantage of nanomaterials to deliver plasmid DNA (*21–25*) or proteins (*26*) to intact plant cells. Polymeric nanoparticles have shown promise for siRNA delivery to cell wall-free plant protoplasts, but polymeric nanoparticles have not been shown to traverse the cell wall for gene silencing in intact plant cells (*13*). A recent study has shown that clay nanosheets can facilitate delivery of pathogen-specific double-stranded RNA into intact plant cells for virus resistance (*27*). Topical application of clay nanosheets enabled silencing of homologous RNA to provide sustained 20-day viral protection on the leaf surface. Clay nanosheet platform is a promising use of nanoparticles for delivery of RNAi into plants, paving the way towards future developments in plant bionanotechnology.

For many applications, particularly biosynthetic pathway mapping, direct and strong but also transient gene silencing is desired within all cellular layers of plant leaves whilst also mitigating against RNA degradation. In this study, we demonstrate the delivery of a different RNAi molecule – single-stranded siRNA – into intact cells of plant leaves using high-aspect-ratio one dimensional carbon nanomaterials: single-walled carbon nanotubes (SWNTs). SWNTs are biocompatible allotropes of carbon that have a high aspect ratio cylindrical nanostructure with diameters of 0.8-1.2 nm and lengths of 500-1000 nm. SWNTs are capable of passively crossing the extracted chloroplast envelope (*28*) and plant cell membranes (*29*) due to their high aspect ratio morphology, uniquely high stiffness, and small dimensions. SWNTs are among the few nanomaterials that can be synthesized to have a smallest dimension (~ 1 nm) below the plant size exclusion limit of ~20 nm, while also providing a large cylindrical surface area from the extrusion of their 1-dimension out to ~ 500 nm. The resulting large surface area to volume ratio is thus amenable to facile loading of appreciable quantities of biological cargoes such as siRNA. In contrast, spherical nanoparticles must often exceed the plant cell wall size exclusion limit to load necessary quantities of bio-cargoes, due to the reduced scaling of the spherical nanoparticle surface area to volume. Furthermore, when bound to SWNTs, biomolecules are protected from degradation in mammalian systems (*30*), exhibiting superior biostability compared to free biomolecules; a phenomenon we show herein can extend to plants. Moreover, SWNTs have strong intrinsic near-infrared (nIR) fluorescence (*31, 32*) within the biological tissue-transparency window and beyond the chlorophyll autofluorescence range, and thus enable tracking of cargo-nanoparticle complexes deep in plant tissues.

Prior usage of SWNTs in plant systems is limited to studies of SWNT biocompatibility (*28, 33, 34*), sensing of small molecules (*29, 35, 36*), and for delivery of plasmid DNA for genetic transformations (*24, 25*). To-date, there has yet to be a nanoparticle-based delivery platform for siRNA molecules into intact plant cells. Herein, we develop a SWNT-based siRNA delivery platform for the efficient silencing of an endogenous *Nicotiana benthamiana* gene in plant leaves. We show that SWNTs enable passive delivery (without external mechanical aid) and fluorescent-tracking of siRNA molecules in plant tissues. SWNTs present a non-toxic platform for siRNA delivery that uses a minimal siRNA dose to achieve silencing for up to 7 days, whereby silencing can be sustained upon re-infiltration of the siRNA-SWNT dose. With SWNT-mediated siRNA delivery, we achieve 95% gene silencing efficiency at the mRNA level, and show a significant delay in siRNA nuclease degradation in cells, and also at the single-molecule level, through protection by SWNTs. Taken altogether, SWNT-based delivery platform is rapid, scalable, facile to multiplex for multiple gene silencing targets, and species-independent (*24, 33, 37–39*). In sum, this study establishes that SWNTs could be a promising resource to overcome plant RNA delivery limitations, and could enable a variety of plant biotechnology applications based on RNAi.

## RESULTS

### Preparation and characterization of siRNA-SWNTs

In this study, we aim to validate SWNTs as a passive and effective siRNA delivery and gene silencing platform for use in intact cells of mature plants. To this end, we aim to silence GFP gene expression in transgenic *mGFP5 Nicotiana benthamiana* (*Nb*) plants by delivering siRNA molecules into leaves with SWNT nanocarriers. *mGFP5 Nb* plants constitutively express GFP targeted to the ER under the control of the *Cauliflower mosaic virus* 35S promoter (*40*) (DNA sequences for the promoter and GFP gene can be found in Supplementary Data 1). Herein, we tested two separate siRNA sequences (a-siRNA and b-siRNA) which target two slightly different regions of the *mGFP5* gene for GFP silencing (Fig. 1a).

**Fig. 1.**
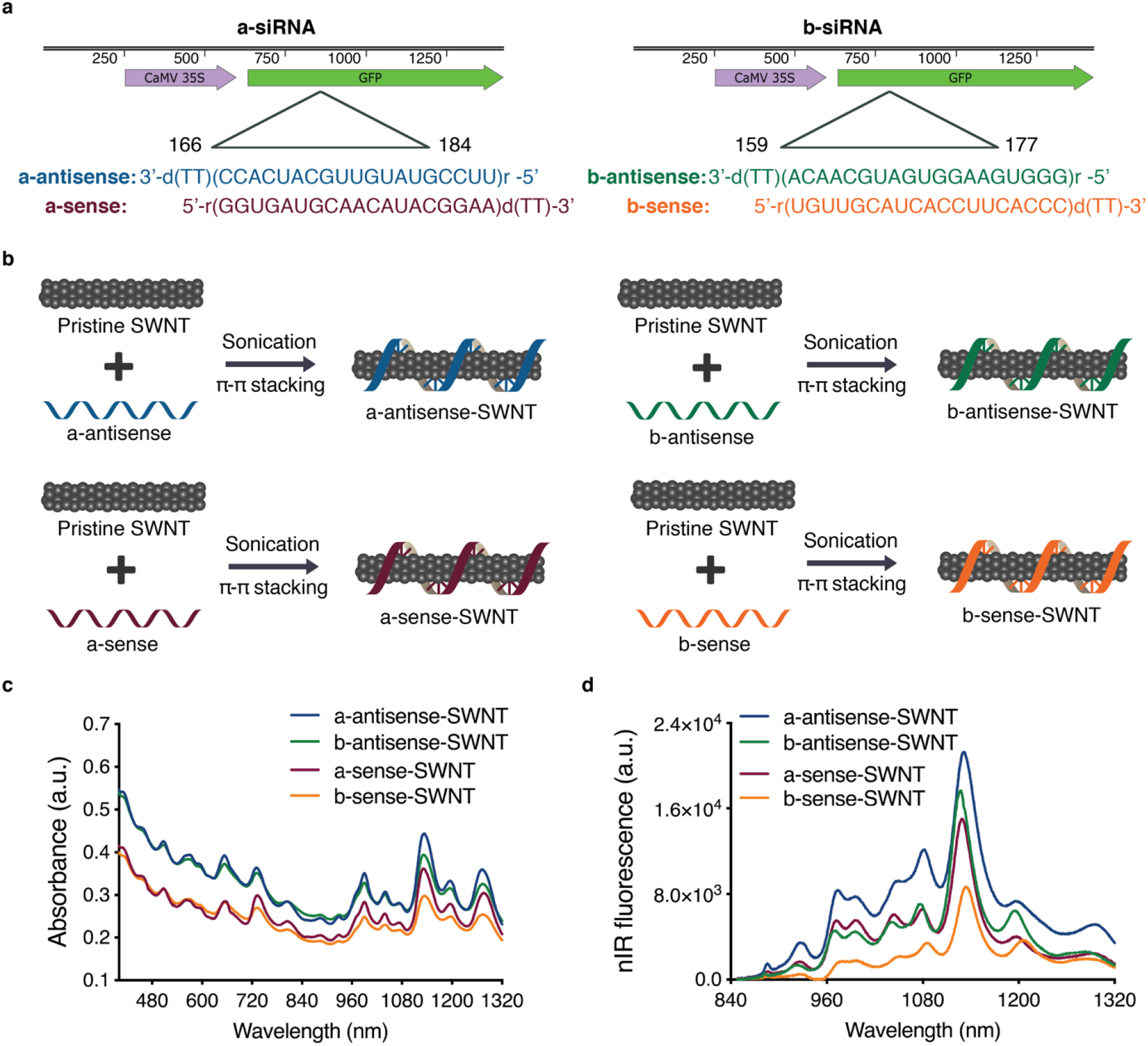
siRNA-SWNT preparation and characterization. **a** Two sets of siRNA sequences targeting the GFP gene of transgenic mGFP5 *Nicotiana benthamiana* were separately tested in this study. Sequences on the left were chosen from Tang *et al*. (*41*) and sequences on the right were designed specifically for this study. **b** Suspension of pristine SWNTs with sense and antisense single-stranded RNA sequences *via* probe-tip sonication. **c** Absorbance spectra of all RNA-SWNT suspensions. **d** nIR spectra of all RNA-SWNT suspensions.

Loading of siRNA on SWNTs was accomplished by probe-tip sonication of each siRNA single-strand (sense, and separately antisense) with pristine SWNTs for both a-siRNA and b-siRNA sequences (Fig. 1b). With this method, sense and antisense strands of siRNA were non-covalently adsorbed on SWNTs *via* π-π stacking of RNA nitrogen bases with the π bonds of sp^2^-hybridized carbons in SWNTs. The adsorption of RNA on SWNTs was confirmed for each sequence (a-antisense-SWNT, a-sense-SWNT, b-antisense-SWNT and b-sense SWNT) through the emergence of characteristic peaks in the individually-suspended SWNT absorbance (Fig. 1c) and nIR fluorescence emission spectra (Fig. 1d). We hypothesize and later verify that upon infiltration of an equimolar mixture of sense and antisense suspended SWNTs, these complementary siRNA strands desorb from the SWNT surface and hybridize to each other inside plant cells to form the active double-stranded siRNA silencing complex.

As a negative control for all siRNA silencing studies, we used SWNTs suspended with a non-targeting scrambled RNA sequence (*42*) (s-RNA-SWNT, Supplementary Table 3), which is not complementary to the *mGFP5* mRNA. Successful suspension of SWNTs with non-targeting RNA sense and antisense strands was confirmed by absorbance and fluorescence spectra of individually suspended s-RNA-SWNTs (Supplementary Fig. 1). Furthermore, the atomic force microscopy (AFM) characterization of single-stranded RNA (ssRNA) suspended SWNTs reveals an average ssRNA-SWNT conjugate length of 776.6 nm and an average conjugate height of 1.567 nm (Supplementary Fig. 1), which agrees with the expected values for undamaged and individually suspended ssRNA-SWNTs.

We first tested the internalization of ssRNA-SWNTs into intact *mGFP5 Nb* leaf cells. All internalization studies were performed with a-antisense-SWNT suspension as a representative strand to demonstrate the internalization ability of single-stranded RNA loaded SWNTs into intact walled plant leaf cells. Cy3 fluorophore-tagged RNA-SWNTs (100 nM siRNA and 2 mg/L SWNTs) and Cy3 tagged free RNA (100 nM) solutions were introduced into the intact plant leaves by infiltrating the abaxial surface of the leaf lamina with a needleless syringe (Fig. 2a). Following 6 hours of incubation, infiltrated *mGFP5 Nb* leaves were imaged with confocal microscopy to quantify Cy3 fluorescence inside leaf cells and in the extracellular area. In plants, the cytosol is pushed to the cell periphery due to the presence of a large central vacuole. Leaves infiltrated with Cy3-RNA-SWNTs showed a high degree of co-localization (70% ± 8%, mean ± SD) between the intracellular (cytosolic) GFP and Cy3 fluorescence originating from the nanocarriers, which confirms efficient internalization of RNA-SWNTs into intact cells (Fig. 2b). Conversely, leaves infiltrated with Cy3-RNA show minimal co-localization between the GFP and Cy3 channels (12% ± 10%, mean ± SD), and Cy3 fluorescence is observed mostly around the guard cells, suggesting free RNA is not able to internalize into intact plant cells efficiently (Fig. 2b). Additional confocal images of Cy3-RNA-SWNT and Cy3-RNA infiltrated leaves with representative higher and lower co-localization percentages are presented in Supplementary Fig. 2. As a note here, a typical plant cell contains an organelle called vacuole, which is filled with water and occupies 80% of the cell volume. Therefore, any fluorescence localized in the cytoplasm follows the cytosolic cell contour shape (Supplementary Fig. 3).

**Fig. 2.**
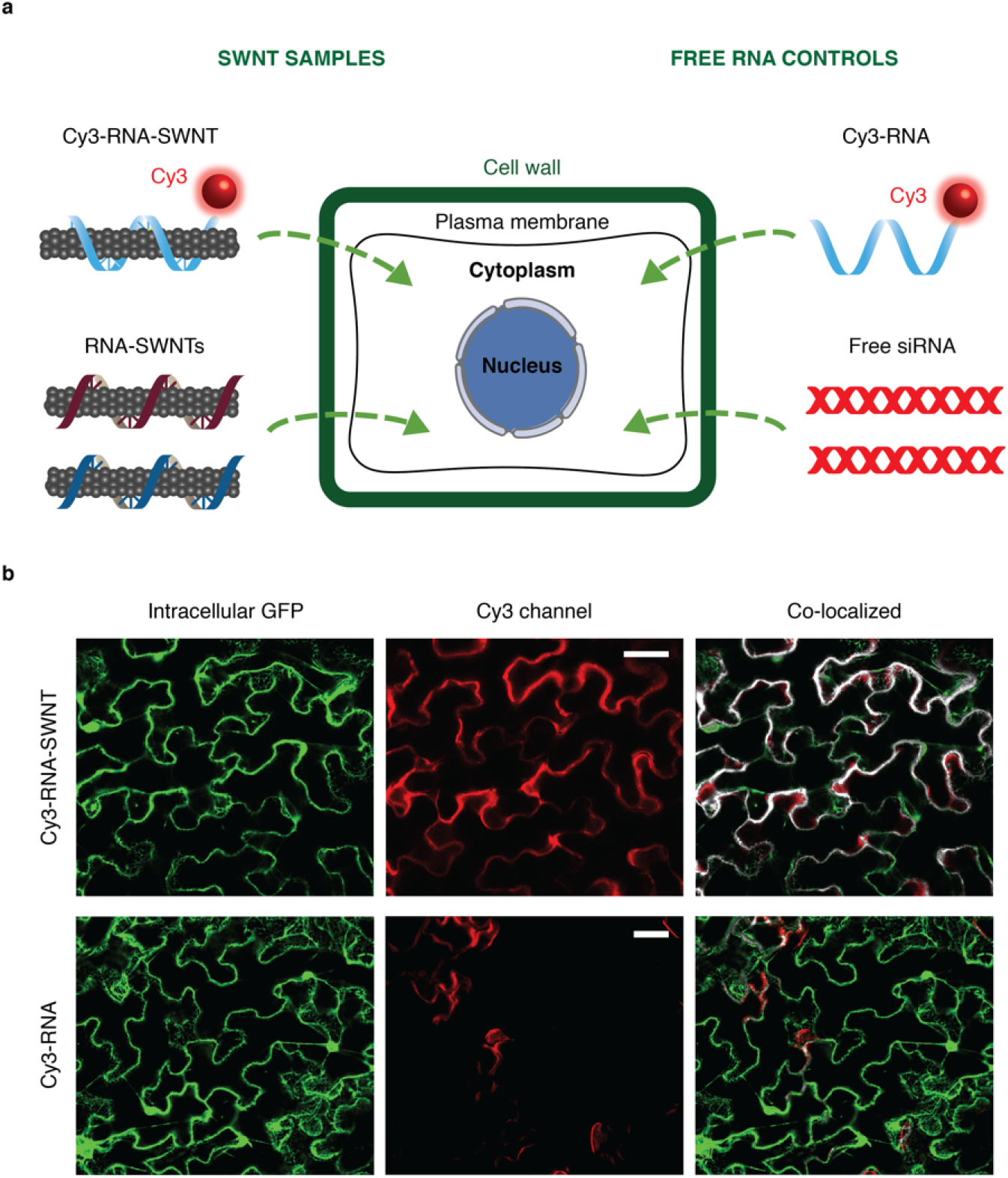
ssRNA-SWNT internalization into transgenic *mGFP5 Nicotiana benthamiana* leaves. **a** Schematic showing samples tested for internalization into *mGFP5 Nb* leaves (Cy3-tagged RNA-SWNTs and Cy3-tagged free RNA as a control), and samples subsequently tested for silencing of a constitutively expressed GFP gene (RNA-SWNTs and free siRNA as a control). **b** Representative confocal images of Cy3-RNA-SWNT and Cy3-RNA infiltrated *Nb* leaves; intracellular GFP (green), Cy3 (red) and co-localization (white) channels. All scale bars are 20 μm.

To investigate the effect of SWNT length on the cell internalization efficiency, we prepared short SWNTs through excessive probe-tip sonication. AFM images revealed that these short SWNTs have an average length of 250 nm; they are significantly shorter than SWNTs obtained with regular preparation (776 nm). We then loaded these short SWNTs with Cy3-RNA as before and checked internalization efficiency into GFP *benthamiana* cells with confocal microscopy. Interestingly, we found that short SWNTs have lower plant cell internalization efficiency compared to the longer ones, shown by respective average co-localization percentages of 47% and 70% (Supplementary Fig. 4).

In addition to confocal imaging of fluorophore tagged ssRNA-SWNTs, we verified internalization of SWNT nanocarriers into intact leaf cells by leveraging the intrinsic SWNT nIR fluorescence. *mGFP5 Nb* leaves were infiltrated with ssRNA-SWNTs or free RNA without a fluorophore (Fig. 2a). Following 6 hours of incubation, we imaged the infiltrated leaves with a custom-built nIR microscope equipped with a Raptor Ninox VIS-SWIR 640 camera, a 721 nm SWNT excitation laser, and a white lamp and appropriate filters to image GFP (see Methods). In leaves infiltrated with ssRNA-SWNTs, commensurate with Cy3-tagged confocal imaging results, we observe a high degree of co-localization between intracellular GFP and the nIR fluorescence of SWNTs (Supplementary Fig. 5), further substantiating efficient internalization of SWNTs into intact plant cells. No co-localization was observed in leaves treated with unlabeled free RNA. The internalization of SWNT nanocarriers into plant cells is also supported by the nIR fluorescence spectra of ssRNA-SWNTs. Compared to as-prepared ssRNA-SWNTs, the nIR fluorescence spectra of ssRNA-SWNTs infiltrated into leaves shows a 6-nm solvatochromic shift, and a relative change in intensity of small bandgap nanotubes upon cell membrane crossing (Supplementary Fig. 5). These differences in SWNT nIR spectra upon infiltration into leaves are possibly the result of the local dielectric environment change and exposure to intracellular biomolecules (*36, 43, 44*).

After confirming that ssRNA adsorbed SWNTs can efficiently be uptaken by plant cells, we analyzed the thermodynamics of sense and antisense strand desorption from the SWNT surface, and their subsequent propensities for hybridization in the extracellular and intracellular conditions. According to our analysis (Supplementary Information), in the *in vitro* and extracellular area of the leaf tissue, sense and antisense strand desorption from the SWNT surface and hybridization is not thermodynamically favorable (ΔG>0), due to a high free energy cost of bare SWNTs in an aqueous environment (Fig. 3a). This unfavorable RNA desorption energy facilitates maintenance of intact RNA-SWNT conjugates in the extracellular environment until RNA-SWNTs enter cells. Once intracellular, sense and antisense strand desorption from the SWNT surface and hybridization is thermodynamically favorable (ΔG<0) because intracellular proteins, lipids, and other membrane and cytosolic biomolecules can occupy the SWNT surface and lower the associated free energy costs of RNA desorption (Fig. 3b).

**Fig. 3.**
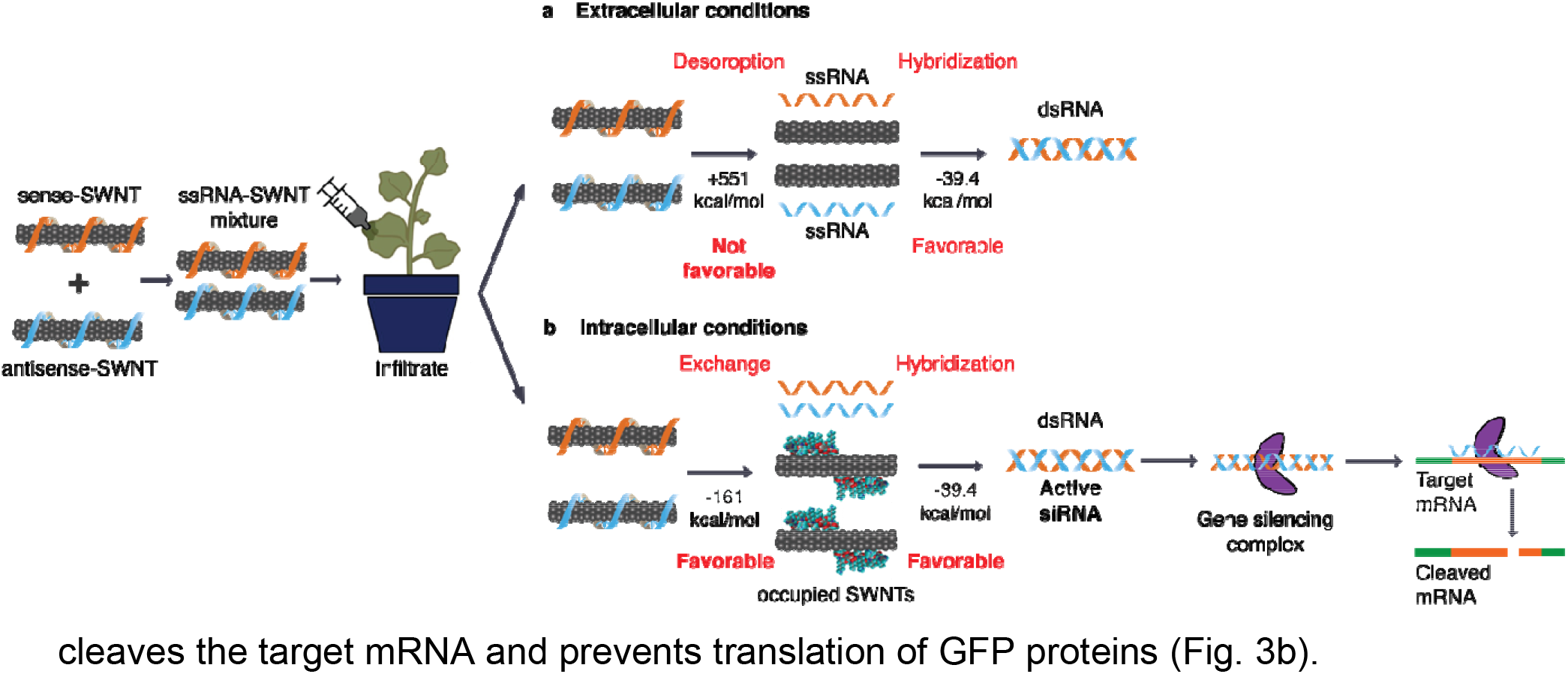
Thermodynamic analysis of RNA desorption from SWNTs and hybridization in extracellular and intracellular conditions, and proposed gene silencing mechanism. **a** An equimolar mixture of sense-SWNT and antisense-SWNT suspensions are infiltrated into transgenic *Nb* leaves with a needleless syringe. In the extracellular area of leaf tissue, RNA desorption and hybridization is not thermodynamically favorable due to the high free energy cost of bare SWNTs. **b** Inside cells, RNA desorption from SWNTs and hybridization is thermodynamically favorable because molecules can occupy the bare SWNT surface and lower the RNA desorption free energy cost. Upon desorption from SWNTs, double-stranded active siRNA assembles with the gene silencing complex and complexes with target mRNA for cleavage and gene silencing.

Hybridization and desorption of sense and antisense RNA strands is verified with an *in vitro* experiment, where we mixed and incubated an equimolar mixture of a-sense-SWNT and a-antisense-SWNT suspensions for 3 h at room temperature; either in water or in plant cell lysate solution (Supplementary Fig. 6). We then eluted the desorbed siRNA and quantified it *via* absorbance at 260 nm. The results confirm that an insignificant amount of siRNA is desorbed when RNA-SWNTs incubated in water, whereas 66% of the siRNA is desorbed when incubated in plant cell lysate solution. We then ran the eluted RNA from the cell lysate sample on an agarose gel and showed that it is double-stranded, which verifies the formation of double-stranded siRNA in the cell cytosol. Additionally, zeta potential measurements of a-siRNA-SWNTs before and after hybridization in water and removal of desorbed RNA show unchanged nanoparticle zeta potential, suggesting there is not significant amount of RNA hybridizing and desorbing from SWNT surface in water (Supplementary Fig. 6).

Once hybridized, double-stranded active siRNA can merge with the gene silencing complex, whereby the antisense strand of siRNA directs the complex to the endogenous target mRNA. Upon hybridization of the antisense strand with the complementary target mRNA, a protein in the gene silencing complex (Argonaute), cleaves the target mRNA and prevents translation of GFP proteins (Fig. 3b).

Following verification of SWNT internalization and formation of active siRNA complexes in plant cells, we next infiltrated transgenic *mGFP5 Nb* leaves with siRNA-SWNTs and control solutions to determine the gene silencing efficiency of this platform. Silencing studies were conducted with the following samples at 100 nM final siRNA and 2 mg/L final SWNT concentration: non-treated leaves, s-RNA-SWNT (non-targeting), free siRNA, a-siRNA-SWNT, and b-siRNA-SWNT (See Supplementary Table 3 for sequences). We have shown that 100 nM siRNA on SWNTs is an optimal dose to be used in *mGFP5* silencing studies (Supplementary Fig. 7). Transgenic *Nb* leaves that constitutively express GFP were imaged *via* confocal microscopy to quantify GFP silencing at the protein level. Representative confocal images of the leaves 2-days post-infiltration reveals that both a-siRNA-SWNTs and b-siRNA-SWNTs lead to significant reduction of GFP in cells, whereas GFP expression in leaves infiltrated with s-RNA-SWNT and free siRNA appears similar to GFP expression in non-treated leaves (Fig. 4a). Quantification of GFP fluorescence intensity from the confocal images of s-RNA-SWNTs and a-siRNA-SWNTs (see Methods) reveals that a-siRNA-SWNT infiltrated leaves have 38% ± 3.2% (mean ± SD) less GFP protein 3-days post-infiltration compared to the s-RNA-SWNT infiltrated leaves. At 7-days post-infiltration, a-siRNA-SWNT shows roughly the same amount of GFP, 106.6 ± 4.1% (mean ± SD), as s-RNA-SWNT infiltrated leaves (Fig. 4b), as expected since gene silencing with siRNA is a transient process. GFP silencing with a-siRNA-SWNT was also verified with a Western blot analysis, where GFP extracted from the *Nb* leaves infiltrated with a-siRNA-SWNT is 42.6% ± 2.8% (mean ± SD) less than GFP extracted from s-RNA-SWNT infiltrated leaves both at 1 and 2-days post-infiltration (Fig. 4c).

**Fig. 4.**
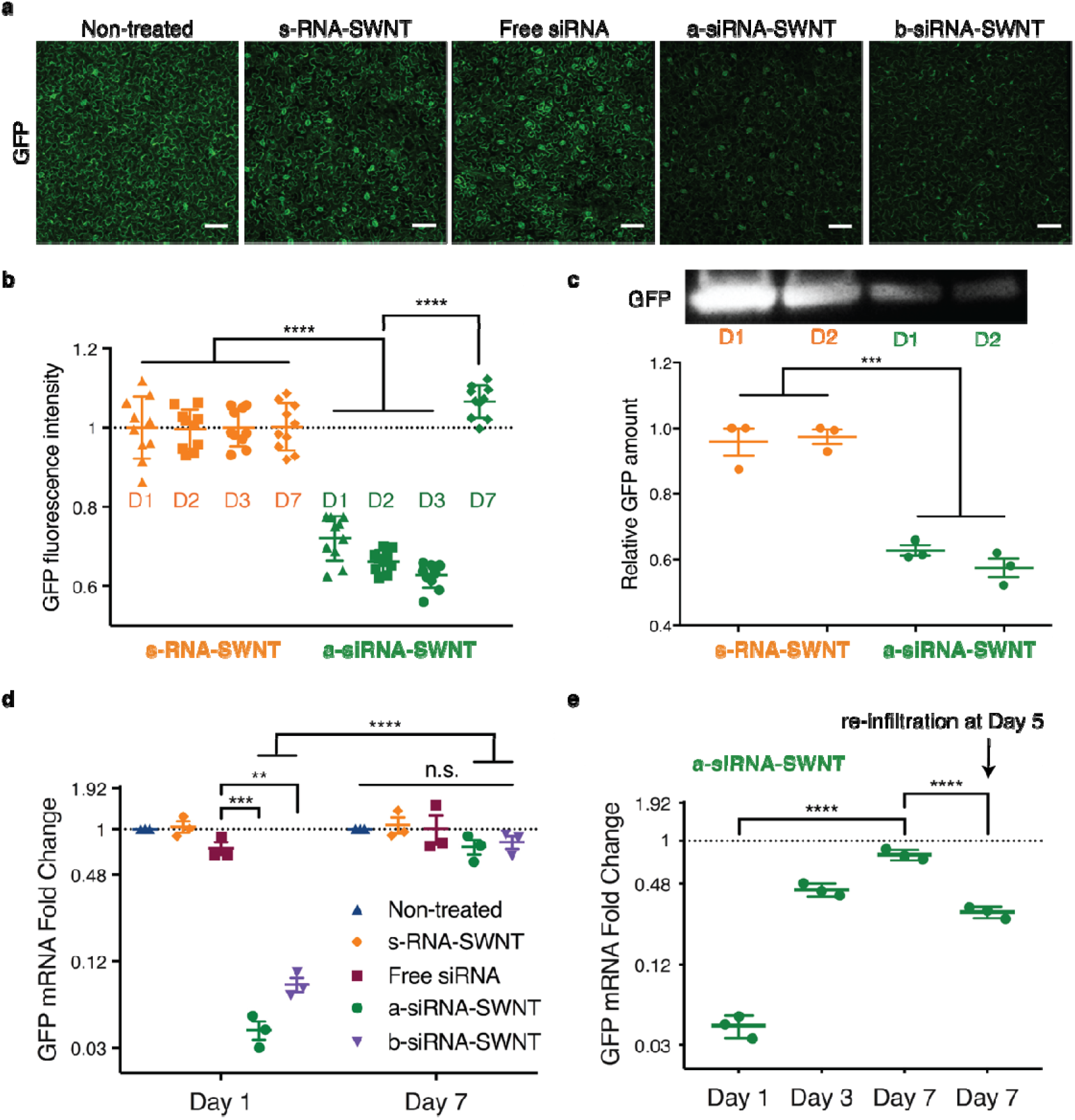
GFP gene silencing with RNA-SWNTs at the mRNA transcript and protein level. **a** Representative confocal microscopy images of non-treated, s-RNA-SWNT, free siRNA, a-siRNA-SWNT, and b-siRNA-SWNT infiltrated transgenic *Nb* leaves 2-days post-infiltration. Scale bars, 100 μm. **b** Quantitative fluorescence intensity analysis of confocal images for s-RNA-SWNT and a-siRNA-SWNT at 1, 2, 3 and 7-days post-infiltration. \*\*\*\**P* < 0.0001 in one-way ANOVA. Error bars indicate s.d. (n = 10). **c** Representative Western blot for GFP extracted from s-RNA-SWNT and a-siRNA-SWNT infiltrated *Nb* leaves 1 and 2 days post-infiltration, and quantification of GFP protein. \*\*\**P* = 0.0001 in one-way ANOVA and error bars indicate s.e.m (n = 3). **d** qPCR analysis for GFP mRNA fold change at Day 1 and 7 post-infiltration for all samples tested. \*\**P* = 0.0016, \*\*\**P* = 0.0008 and \*\*\*\**P* < 0.0001 in two-way ANOVA (n.s: non-significant) All error bars indicate s.e.m. (n = 3). **e** qPCR analysis for GFP mRNA fold change at Day 1, 3, 7 and Day 7 with re-infiltration at Day 5 for a-siRNA-SWNT treated *Nb* leaf sample. \*\*\*\**P* < 0.0001 in one-way ANOVA and all error bars indicate s.e.m. (n = 3). All qPCR data for GFP expression are normalized with respect to housekeeping gene Elongation Factor 1 (EF1), and a control sample of a non-treated leaf.

We corroborated the GFP reduction results obtained with confocal imaging and Western blot analysis by performing quantitative PCR (qPCR) at the mRNA transcript level. One day after infiltration of leaves with s-RNA-SWNT, free siRNA, a-siRNA-SWNT, and b-siRNA-SWNT, we extracted total RNA from the leaves and quantified the GFP mRNA transcript levels in each sample at Day 1 and 7. qPCR demonstrates that s-RNA-SWNT and free siRNA infiltrated leaves have the same amount of GFP mRNA transcript as the non-treated leaf, whereby a-siRNA-SWNT and b-siRNA-SWNT infiltrated leaves show 95% ± 4.1% (mean ± SD) and 92% ± 6.2% (mean ± SD) reduction in the GFP mRNA transcript levels at Day 1, respectively (Fig. 4d). Similar to the confocal results, we found that mRNA transcript levels return back to the baseline levels as observed in non-treated leaves by Day 7 in all samples as a result of transient silencing (Fig. 4d). Additionally, we show that we can recover GFP silencing at Day 7 by up to 71% ± 2.9% (mean ± SD) by re-infiltrating the leaf with second 100 nM a-siRNA-SWNT dose at Day 5 (Fig. 4e). With the same technique, we also demonstrated the silencing of a functional endogenous *Nicotiana benthamiana* gene called ROQ1, which has implications in disease resistance against many pathogens (*45*) (Supplementary Fig. 8). Our results verify that SWNTs can also silence endogenous plant genes efficiently.

It is likely that SWNT scaffolding improves internalization of siRNA and also protects siRNA from degradation once intracellular. To explore this hypothesis, we performed single molecule total internal reflection fluorescence (smTIRF) microscopy to probe single siRNA strand susceptibility to degradation by RNase A when adsorbed on SWNTs, compared to single free siRNA. To do so, we labeled the a-antisense strand of GFP siRNA with a 5’ terminal Cy3 fluorophore, and immobilized RNA-Cy3 and RNA-Cy3-SWNTs onto parallel channels of a microfluidic slide (see Methods). We measured the Cy3 fluorescence in each channel before and after treatment with RNase A, whereby percent decrease in the number of Cy3 molecules was used as a proxy for the percent siRNA degraded (Fig. 5a). Our TIRF results show that 98% ± 0.3% (mean ± SD) of the initial Cy3-RNA immobilized on the channel surface is degraded after incubation with RNase A, whereas only 16% ± 4.9% (mean ± SD) of Cy3-RNA is degraded when it is bound to SWNTs, suggesting that SWNTs protect the siRNA cargo from enzymatic degradation inside cells (Fig. 5b). Negative controls in which only salt buffer is flown through, or empty BSA-passivated channels, do not show appreciable changes in fluorescence or fluorescence counts, respectively (Supplementary Fig.9).

Intracellular stability of single stranded RNA (ssRNA) suspended SWNTs and free ssRNA was also assessed by incubating ssRNA-SWNT conjugates with total proteins extracted from plant leaves (*i.e.* plant cell lysate). Agarose gel electrophoresis of free ssRNA *vs*. ssRNA-SWNTs incubated in plant cell lysate for 1, 3, 6, 12, and 24 hours demonstrate that free ssRNA is degraded significantly faster in cells compared to ssRNA adsorbed on SWNTs (Fig. 5c). Band intensity quantification of agarose gel reveals that upon starting with 200 ng ssRNA, free ssRNA is completely degraded within 12 hours, whereas the ssRNA on the SWNTs is only completely degraded after 24 hours (Fig. 5d and 5e), which corresponds to a 12 hour increase in the residence time of siRNA strands in cells when delivered through SWNTs. This gives rise to prolonged and increased silencing efficiency, as siRNA strands in cells have higher chance of hybridizing into the active complex before getting degraded by plant nucleases. With a similar *in vitro* cell lysate degradation experiment, we also show that after hybridization and desorption, double-stranded siRNA has high stability and it persists in cells for more than 4-days after formulation (Supplementary Fig. 10).

**Fig. 5.**
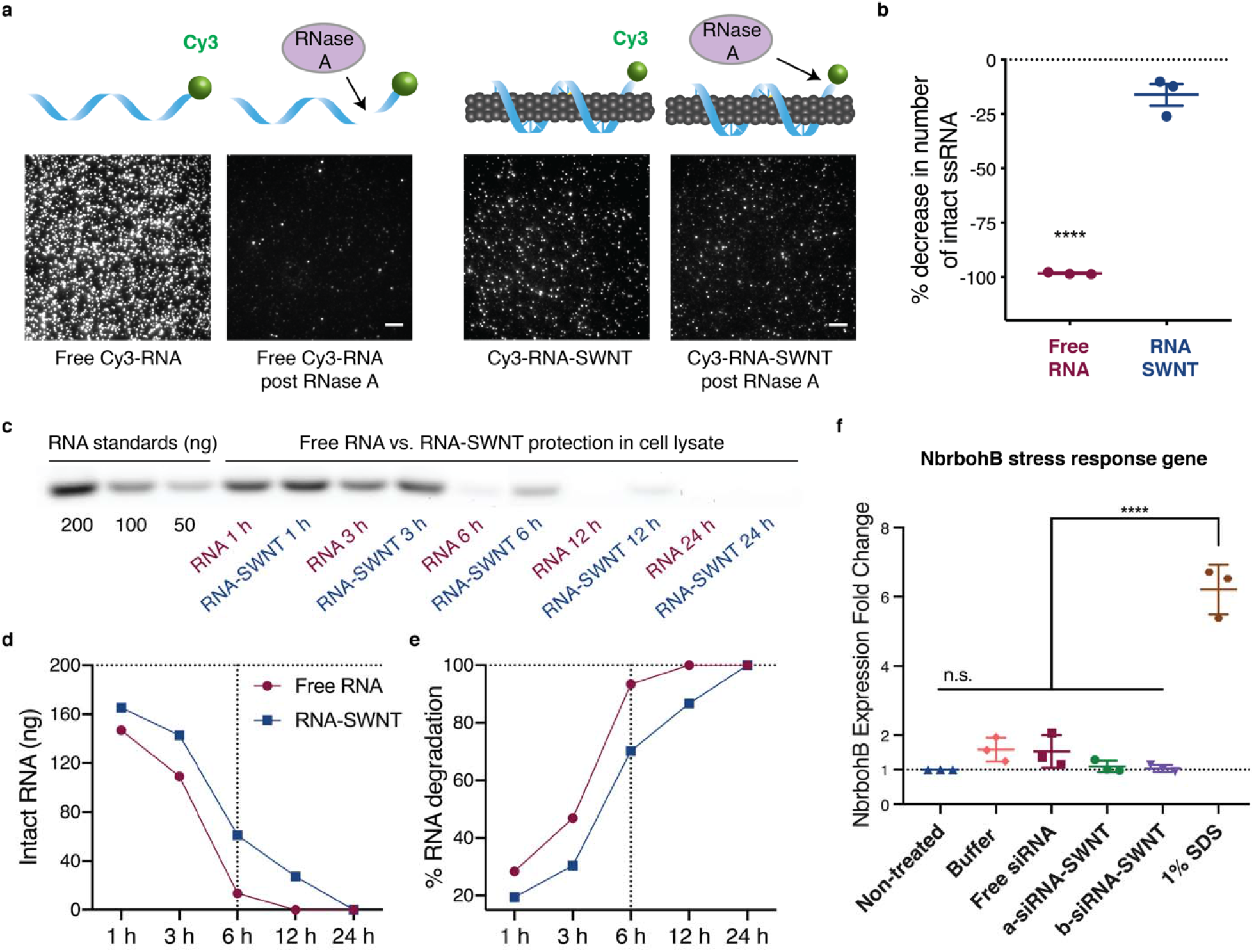
RNA protection from enzymatic degradation and SWNT toxicity analysis. **a** smTIRF microscopy of Cy3-labeled RNA and Cy3-labeled RNA-SWNTs before and after incubation with RNase A. Scale bars, 5 μm. **b** Quantification of % decrease in number of intact RNA molecules upon RNase A treatment. Error bars indicate s.e.m. (n = 3). \*\*\*\**P* < 0.0001 in two-tailed unpaired t-test. **c** Agarose gel electrophoresis of free RNA and RNA-SWNTs incubated in plant cell lysate for 1, 3, 6, 12, and 24 hours. **d** Quantification of intact RNA from the agarose gel in part c. **e** Quantification of % RNA degradation from the agarose gel in part c. **f** qPCR analysis of *NbrbohB* following a 3-hour exposure to samples. \*\*\*\**P* < 0.0001 in one-way ANOVA and error bars indicate s.e.m. (n = 3).

SWNT biocompatibility, at the concentrations used in this study, was tested by qPCR analysis of a commonly used stress gene, and by tissue damage analysis *via* confocal microscopy. For qPCR toxicity analysis, we checked the upregulation of the respiratory burst oxidase homolog B (*NbrbohB*) gene (Fig. 5f). *NbrbohB* upregulation in *Nicotiana benthamiana* leaves represents stress response to many factors such as mechanical, light or heat damage (*46*). qPCR results show that 2 mg/L RNA-SWNT treated areas in leaves do not upregulate *NbrbohB* gene compared to buffer treated adjacent areas within the same leaves. 1% SDS solution was used as a positive toxicity control, and upregulated *NbrbohB* gene by 6-fold 3 hours post-infiltration (Fig. 5f). Tissue damage in the RNA-SWNT and 1% SDS infiltrated leaves was also monitored *via* confocal microscopy, and no tissue or cell damage was detected in RNA-SWNT infiltrated leaves, whereas significant distortion of cell morphology and tissue integrity can be seen in the SDS treated areas (Supplementary Fig. 11). Given the unchanged expression levels of stress gene *NbrbohB,* and healthy leaf tissue of RNA-SWNT infiltrated plants, we can conclude that 2 mg/L RNA-SWNTs are biocompatible for *in planta* RNAi applications.

## DISCUSSION

Nanomaterials have shown much promise for plasmid (*24, 25*) and protein (*26*) delivery to plants, motivating their use for plant delivery of RNAi, as has proven quite fruitful for human therapeutics. We demonstrate here that high-aspect-ratio one dimensional SWNTs can successfully deliver siRNA molecules to efficiently silence a GFP gene in transgenic *Nicotiana benthamiana* mature plant leaves, through a combination of i) effective intracellular delivery and ii) protection of the siRNA cargo from nuclease degradation. We found that RNA adsorbed SWNTs rapidly and efficiently internalize into the full leaf thickness of mature walled plant cells within 6 hours, in contrast to free RNA internalization which is minimal. We further found that π-π adsorption of siRNA on the SWNT surface delays intracellular siRNA degradation and thus prolongs silencing.

Here, we developed a platform for siRNA delivery using nanoparticles, well suited for cellular delivery in plant tissues with intact cell walls. This platform utilizes SWNTs, to which single-stranded sense and antisense siRNA are adsorbed separately, enabling thermodynamically-favorable siRNA hybridization once intracellular for subsequent gene silencing mechanisms. We show that ssRNA is protected from degradation for up to 24 hours when adsorbed to SWNTs, whereas free ssRNA is almost completely degraded by 6 hours. We show a similar siRNA protection phenomenon with single-molecule TIRF microscopy of individual siRNA molecules either free or adsorbed to SWNTs. With this rapid and facile SWNT delivery platform, we achieve transient and DNA-free silencing of genes in mature plant leaves with a low siRNA-SWNT dose, showing mRNA knockdown efficiencies of up to 95% within 1 day post-infiltration, returning to native transcript levels by day 7. We further show that it is possible to retain gene silencing for longer periods of time with a re-infiltration of another siRNA-SWNT dose at day 5, for applications in which sustained silencing is desired. Applications that require the introduction of repeated doses of siRNA-SWNTs may cause some long-term toxicity due to the nanoparticle accumulation in cells. However, studies should be undertaken to investigate the long-term effects of SWNT accumulation in plant cells.

The commonly used cationic nanoparticles for the delivery of negatively charged siRNA through electrostatic interactions have shown appreciable cellular toxicity to cells for certain effective concentrations and/or charge densities (*47*). The pristine non-charged SWNT surface eliminates this problem and makes it possible to scale-up the delivery of siRNA for higher doses or systemic administration. Additionally, the platform could be adapted to loading multiple siRNA sequences to multiplex gene silencing targets by delivering a mixture of SWNTs suspended with multiple siRNA sequences or loading a single SWNT sample with multiple siRNA sequences. Furthermore, SWNT internalization and polynucleotide delivery into plants is hypothesized to be species-independent, can be used with monocots, non-model species, hard-to-transform species, and cargo-carrying SWNTs are expected to diffuse into the full thickness of leaves providing a uniform transformation profile (*24*).

Given aforementioned advantages, we believe that there is a broad range of applications of our siRNA delivery platform. The process of RNA adsorption to SWNTs is based on π-π adsorption and thus agnostic to the function of the RNA cargo. Additional to the more traditional applications of RNAi in plants, such as disease/virus resistance, discovery of biosynthetic pathways, increasing the yield of small-molecule production, and understanding protein functions, SWNT-mediated gene silencing could also potentially be used for efficient and DNA-free delivery of other synthetic ribonucleic acids. For instance, SWNTs could aid nuclease-based genome editing in plants by delivery of single guide RNAs (sgRNAs) and/or messenger RNAs (mRNAs) for controlled and transient nuclease expression and subsequent genome editing. Another potential application of SWNT-based RNA delivery is for increasing homology-directed repair (HDR) rates in plants for gene knock-in applications, which could possibly be achieved by suppressing the expression of the genes required for competitive non-homologous end joining (NHEJ) repair pathways (*48*). As the efficient suppression of these genes is only desirable for the few-day time window in which genome editing takes place, our SWNT-mediated gene silencing platform could enable such control over transient siRNA delivery. As such, SWNT-based delivery of polynucleic acids is a useful resource to expand the plant biotechnology toolkit.

## MATERIALS AND METHODS

### Preparation of chemicals

Super purified HiPCO SWNTs (Lot # HS28-037) were purchased from NanoIntegris, and SWNTs samples were extensively purified before use (*49*). Single-stranded RNA strands, Cy3-tagged single-stranded RNA strands and all primer sequences were purchased from IDT and dissolved in 0.1M NaCl before use. 100K MWCO Amicon spin filters were purchased from Fisher Scientific. The following chemicals were purchased from Sigma-Aldrich: sodium dodecyl sulfate (molecular biology grade), sodium chloride, Tris/HCl, EDTA, NP-40, glycerol, BSA-Biotin and NeutrAvidin. RNAse A was purchased from Takara Bio. All PCR reagents and materials, and molecular biology grade agarose were purchased from Bio-Rad. UltraPure DNase/RNase-free distilled water from Invitrogen was used for qPCR, and EMD Millipore Milli-Q water was used for all other experiments.

### Plant growth

Transgenic mGFP5 *Nicotiana benthamiana* seeds were kindly provided by the Staskawicz Lab, UC Berkeley. The seeds were germinated and grown in SunGro Sunshine LC1 Grower soil mix for four to six weeks before experiments in a growth chamber, 12-hour light at 24°C and 12-hour dark at 18°C cycle. All experiments were done with intact leaves attached to plants, where plants were incubated in the growth chamber until the time of data collection.

### RNA-SWNT and Cy3-RNA-SWNT preparation

SWNTs were suspended with single-stranded RNA polymers or Cy3-tagged single-stranded RNA sequences through probe-tip sonication as previously described (*50*). See **Supplementary Table 3** for all RNA sequences used in this study. Briefly, RNA was dissolved in 0.1 M NaCl at a concentration of 100 mg/mL. 1 mg dry HiPCO SWNTs was added to 20 μL of dissolved RNA, and the solution volume was completed to 1 mL with 0.1 M NaCl. The mixture of SWNTs and RNA was bath sonicated for 10 min at room temperature. Then it was probe-tip sonicated with a 3-mm tip at 50% amplitude (~7W) for 30 min in an ice bath. The sonicated solution incubated at room temperature for 30 minutes and centrifuged at 16,100g for 1 h to remove bundled SWNT and any leftover metal catalyst precursor from SWNT synthesis. Any RNA that was not bound to SWNTs was removed *via* spin-filtering 8 times with 100K Amicon filters, and the SWNT concentration of RNA-SWNTs was determined by measuring the carbon nanotube absorbance at 632 nm. Absorbance spectra of RNA-SWNTs were collected with Shimadzu UV-3600 Plus, and fluorescence spectra of RNA-SWNTs were collected with a near-infrared spectrometer (Princeton Instruments IsoPlane 320 coupled to a liquid nitrogen-cooled Princeton Instruments PyLoN-IR 1D array of InGaAs pixels). RNA concentration on suspended SWNTs was determined by measuring the amount of RNA in flow-through solutions after each spin-filter step *via* absorbance at 260 nm, and subtracting the total amount of free RNA washed from the total amount of RNA added.

In more detail, for each suspension, we start with 1 mg of SWNTs and 2 mg of RNA in 1 mL 0.1 M NaCl solution. After the probe-tip sonication and centrifugation, we end up with approximately 40 μg/mL SWNTs, meaning that our SWNT yield is 40 μg/1000 μg = 4%. In terms of siRNA yield, after the probe-tip sonication, centrifugation and removal of free RNA, we end up with 640 μg/mL RNA on SWNTs, meaning that our RNA yield is 640 μg/2000 μg = 32%. These values can slightly change from experiment to experiment, therefore, we made sure to use the same final diluted concentration of siRNA-SWNTs for every experiment at 100 nM siRNA and 2 mg/L SWNT.

### AFM characterization

AFM characterization of RNA-SWNTs was performed as described in (*24*).

### Infiltration of leaves with RNA-SWNTs and control solutions

Healthy and fully-developed leaves from mGFP5 *Nicotiana benthamiana* (4-6 weeks old) plants were selected for experiments. A small puncture on the abaxial (bottom) surface of the leaf was introduced with a pipette tip, and ~100 μL of the RNA-SWNT solution was infiltrated from the hole with a 1 mL needleless syringe with caution not to damage the leaf.

### Internalization imaging with confocal and nIR fluorescence microscopy

The a-antisense siRNA strand was utilized in the internalization study. After infiltration of 100 nM RNA carrying 2 mg/L RNA-SWNTs, plants with attached infiltrated leaves were left in the plant growth chamber to allow for internalization for 6 h, and imaged with confocal microscopy to track Cy3-tagged RNA-SWNTs in leaves. A Zeiss LSM 710 confocal microscope was used to image the plant tissue with 488 nm laser excitation with a eGFP filter cube to detect intracellular GFP, and 543 nm laser excitation with an appropriate filter cube to detect Cy3 fluorescence. The emission window of Cy3 was adjusted to 550-600 nm to avoid crosstalk between Cy3 and leaf chlorophyll autofluorescence. For nIR imaging, 40 mg/L RNA-SWNTs were infiltrated into leaves and plants with attached infiltrated leaves were left in the plant growth chamber to allow for internalization for 6 h, and imaged with nIR microscopy to track intrinsic SWNT nIR fluorescence in leaves. RNA-SWNT leaf internalization images were captured using a custom-built microscope equipped with a Raptor Ninox VIS-SWIR 640 camera. A 1050-nm long pass filter was used to avoid chlorophyll autofluorescence, and a white lamp with an appropriate filter cube was used to image GFP. GFP and Cy3 (or nIR) images were analyzed with the ImageJ co-localization program to demonstrate internalization of RNA-SWNTs into cells.

### *In vitro* RNA hybridization and desorption assay

a-sense-SWNT and a-antisense-SWNT solutions were prepared according to “**RNA-SWNT and Cy3-RNA-SWNT preparation**” section. Equimolar mixtures of these suspensions each containing 600 ng/μL RNA on SWNTs were either incubated in water or in plant cell lysate for 3 h at room temperature to allow for hybridization and desorption. Next, hybridized double-stranded RNA in solution was eluted with 100K spin filters and the concentration of RNA in the elute was measured *via* absorbance at 260 nm with Nanodrop. For zeta potential measurements in Supplementary Fig. 4c, an equimolar mixture of a-sense-SWNT and a-antisense-SWNT suspensions were incubated in water for 3 h at room temperature to allow for hybridization and desorption. Next, hybridized double-stranded RNA in solution (if any) was eluted with 100K spin filters and the zeta potential of the remaining RNA-SWNT mixture was measured with Malvern Zetasizer.

### Confocal imaging for silencing and quantitative fluorescence intensity analysis of GFP expression

mGFP5 *Nb* leaves were infiltrated with s-RNA-SWNT, free siRNA, a-siRNA-SWNT, and b-siRNA-SWNT at the same RNA concentration of 100 nM and SWNT concentration of 2 mg/L. Infiltrated plant leaves were prepared for confocal imaging 1, 2, 3, and 7-days post-infiltration as described in (*24*). For each sample, mean fluorescence intensity value was normalized with respect to the mean GFP fluorescence intensity of a non-treated leaf. The same imaging parameters and quantification analyses were applied to samples imaged on different days.

### Quantitative Western blot experiments and data analysis

Whole leaves fully infiltrated with samples were harvested 24 and 48 h post-infiltration, and total proteins were extracted as described in (*24*). After quantification of the total extracted proteins by a Pierce 660 nm Protein Assay (Thermo, Prod# 22660), 0.5 μg of normalized total proteins from each sample were analyzed by 12% SDS–PAGE and blotted to a PVDF membrane. The membrane was blocked for 1 hour using 7.5% BSA in PBST (PBS containing 0.1% Tween20) buffer and incubated overnight at 4°C with the primary GFP antibody as required (1:2000 dilution, Abcam, ab290). After extensive washing, the corresponding protein bands were probed with a goat anti-rabbit horseradish peroxidase-conjugated antibody (1:5000 dilution, Abcam, ab205718) for 30 min. The membrane was then developed by incubation with chemiluminescence (Amersham ECL prime kit) plus and imaged by ChemiDoc™ XRS+ System. The intensity of GFP bands were quantified with ImageJ software.

### Quantitative PCR (qPCR) experiments for gene silencing

Two-step qPCR was performed to quantify GFP gene silencing in transgenic *Nb* plants as described in (*24*). The target gene in our qPCR was *mGFP5* (GFP transgene inserted into *Nb*), and *EF1* (elongation factor 1) as our housekeeping (reference) gene. Primers for these genes can be found in **Supplementary Table 3.** An annealing temperature of 60°C and 40 cycles were used for qPCR. qPCR data was analyzed by the ddCt method (*51*) as described in (*24*). For each sample, qPCR was performed as 3 reactions from the same isolated RNA batch, and the entire experiment consisting of independent infiltrations and RNA extractions from different plants was repeated 3 times (3 biological replicates).

### Single molecule TIRF to image RNA protection by SWNTs

The a-antisense siRNA strand was utilized in this assay. 10 μM 5’ labelled Cy3-RNA was added to an equal mass of SWNTs. The RNA-SWNT suspension and removal of unbound RNA followed the same protocol as described in ‘Quantitative PCR (qPCR) experiments for gene silencing’. The positive control comprised of the same sequence that was 5’ Cy3 labeled, and 3’ biotin labeled. 6-channel μ-slides (ibidi, μ-Slide VI 0.5 Glass Bottom) were initially washed by pipetting 100 μL of 100 mM sterile NaCl solutions into one reservoir and removing 60 μL the other end, leaving just enough solution to fully wet the channel. Each subsequent step involved depositing the desired solution volume into the reservoir and removing the equivalent volume from the other end of the channel. Slide preparation was done as described by Kruss and Landry *et al. (52)* with some modifications. Briefly, 50 μL of 0.25 mg/mL BSA-Biotin was added to coat the surface of the glass slide for 5 minutes. Next, 50 μL of 0.05 mg/mL NeutrAvidin was added, followed by 50 μL of 1.0 mg/L RNA-SWNT, which non-specifically adsorbs to NeutrAvidin. For the positive control, 50 μL of 200 pM biotinylated Cy3-RNA was added in place of RNA-SWNT. The addition of each component comprised of a 5-minute incubation period, followed by flushing the channel with 50 μL of NaCl solution to remove specimens that were not surface-immobilized. Each channel was exposed to 50 μL of 10 μg/mL RNase A for 15 minutes at room temperature. The reaction was stopped by rinsing the channel with 50 μL NaCl solution. Slides were imaged with a Zeiss ELYRA PS.1 microscope immediately following incubation with RNase A.

### RNA protection gel assay

To determine if SWNT adsorbed RNA is protected from nuclease degradation, we performed an agarose gel electrophoresis based RNA protection assay as described in (*24*). 200 ng free RNA and RNA-SWNTs (carrying 200 ng RNA) were each incubated with cell lysate proteins obtained from one *Nb* leaf to mimic the intracellular degradation conditions for 1, 3, 6, 12, and 24 hours. After incubation in cell lysate, all RNA (intact or not) was desorbed from the SWNT surface by heating at 95°C for 1 hour in 2% SDS and 1.5 M NaCl solution. Desorbed RNA and cell lysate treated free RNA were run on a 1% agarose gel with RNA standards (200, 100, and 50 ng) to measure the intact *versus* degraded RNA in each sample lane. RNA amounts on the agarose gel were quantified by using band intensity as a proxy (ImageJ Gel Analyzer) and normalized with the lanes containing known RNA quantities.

### dsRNA degradation gel assay

a-sense and a-antisense siRNA strands were hybridized by heating at 95°C for 5 min and 37°C for 1 hour. Hybridized double stranded siRNA samples were incubated in nuclease-free water and cell lysate solutions at room temperature for 16, 24, 48, 72 and 96 hours, and solutions were run on 2% agarose gel. Quantification of the RNA bands from the gel was done using Image J gel analyzer tool. All band intensities were normalized with respect to the hybridized RNA band intensity at time zero without any treatment.

### Plant toxicity analysis

qPCR was used to determine the expression levels of an oxidative stress gene (*NbRbohB*)(*46*) in *Nicotiana benthamiana* plants treated with RNA-SWNTs and control solutions (primer sequences in **Supplementary Table 3**). The samples tested for toxicity were: buffer (0.1 M NaCl), 100 nM free siRNA, a-siRNA-SWNT, b-siRNA-SWNT (each containing 100 nM siRNA and 2 mg/L SWNT) and 1% SDS (as a positive toxicity control), and the qPCR was performed 3-hours after the infiltration of these samples. *EF1* gene was used as a housekeeping gene with an annealing temperature of 60°C for 40 cycles. Same ddCt method was used to analyze the qPCR data (*24*).

### Statistics and data analysis

#### GFP silencing confocal data

N = 10 technical replicates (10 different fields of view from the same leaf per sample infiltration) were imaged. Confocal images reported in Figure 4a are representative images chosen from 10 replicates of Day 2 data. Data are expressed as each mean from the 10 replicates together with error bars indicating standard deviation. Significance is measured with one-way ANOVA with Tukey’s multiple comparisons test. In Figure 4b, F = 124.3 and *P* < 0.0001.

#### Western blot experiment

N = 3 replicates are independent experiments, and Figure 4c denotes the results from a representative blot. Relative GFP amount data determined from the Western blot are expressed as mean from the 3-independent experiments together with error bars indicating standard error of the mean. Significance is measured with one-way ANOVA with Tukey’s multiple comparisons test. F = 54.65, s-RNA-SWNT *vs*. a-siRNA-SWNT *P* = 0.0001.

#### qPCR experiments

For GFP mRNA fold change experiments in Figure 4d, N = 3 replicates are independent experiments, starting with RNA extraction from different leaves through the qPCR amplifications. Each qPCR reaction in 3 independent experiments is performed in triplicate. GFP mRNA fold change data are expressed as each mean from the 3-independent experiments together with error bars indicating standard error of the mean. Significance is measured with two-way ANOVA with Sidak’s multiple comparisons test. Free siRNA *vs*. a-siRNA-SWNT *P* = 0.0008, Free siRNA *vs*. b-siRNA-SWNT *P* = 0.0016, and siRNA-SWNT Day 1 *vs*. Day 7 *P* < 0.0001.

For qPCR results reported in Figure 4e, N = 3 replicates are independent experiments; 3 separate leaves infiltrated per sample and each measured with qPCR. Each sample in each independent experiment consisted of 3 technical replicates of the qPCR reaction. Data are expressed as each mean from the 3-independent experiments together with error bars indicating standard error of the mean. Significance is measured with one-way ANOVA with Tukey’s multiple comparisons test. F = 143.7, Day 1 *vs*. Day 7 *P* = < 0.0001, and Day 7 *vs*. Day 7 (re-inf. at Day 5) *P* = < 0.0001.

#### smTIRF microscopy data

For each sample, N = 3 replicates are 3 channels on a microfluidic slide that were prepared independently. Each channel was imaged to obtain 30 fields of views (technical replicates). In Figure 5b, data are expressed as each mean from the 3-independent channels together with error bars indicating standard error of the mean. Significance is measured with a two-tailed unpaired t-test. F = 317.6 and *P* < 0.0001.

#### Toxicity qPCR data

N = 3 replicates are independent experiments with separate infiltrations of SWNT solutions for each replicate. For the toxicity plot in Figure 5f, 1% SDS *vs*. all other samples *P* < 0.0001 in one-way ANOVA with Tukey’s multiple comparisons test, F = 82.95.

## Data Availability

The DNA sequence of the GFP gene silenced in this study is added as Supplementary Data 1 file in the FASTA format. The data that support the plots within this paper and other findings of this study are available from the corresponding author upon reasonable request.

Supplementary material for this article is available at xxx

The Supporting Information contains the discussion of “Thermodynamic analysis of RNA desorption from SWNTs and hybridization”, and Figures S1–S11, Table S1 and S2 for thermodynamics calculations and Table S3 for RNA sequences and primers used in this study.

## Acknowledgements

We acknowledge support of a Burroughs Wellcome Fund Career Award at the Scientific Interface (CASI), a Stanley Fahn PDF Junior Faculty Grant with Award # PF-JFA-1760, a Beckman Foundation Young Investigator Award, a USDA AFRI award, a USDA NIFA award, and an FFAR New Innovator Award (M.P.L). M.P.L. is a Chan-Zuckerberg Biohub investigator. G.S.D. is supported by a Schlumberger Foundation Faculty for the Future Fellowship. We acknowledge the support of UC Berkeley Molecular Imaging Center, the QB3 Shared Stem Cell Facility, and the Innovative Genomics Institute (IGI).

## Author contributions

G.S.D. and M.P.L. conceived of the project, designed the study, and wrote the manuscript. G.S.D. performed the majority of experiments and data analysis. H.Z. performed Western blot studies and helped with qPCR experiments. N.S.G. performed TIRF experiments and analyzed TIRF data. R.C. prepared some of the RNA-SWNT suspensions used in the studies. All authors have edited and commented on the manuscript, and have given their approval of the final version.

**Correspondence and requests for materials** should be addressed to M.P.L.

## Competing interests

The authors declare no competing interests.

## Notes

#### Summary of Updates

Revised thermodynamic calculations and updated author list

