## Supplemental Information for "Carbon Nanocarriers Deliver siRNA to Intact Plant Cells for Efficient Gene Knockdown"

**Thermodynamic analysis of RNA desorption and hybridization**

Herein, we perform a thermodynamic analysis to model whether siRNA sense and antisense complementary strands desorb from the SWNT surface and hybridize to each other in either the intracellular or extracellular environment (**Fig. 3**). In both cases, it is assumed that the ssRNA desorption from the SWNT surface is a first step, which is followed by hybridization of free ssRNA to dsRNA (nucleic acid binding to SWNTs is through π-π interactions and H-bonding to neighboring strands, therefore, in order to hybridize and form Watson-Crick base-pairing H-bonds, the RNA strands first need to desorb from the SWNT surface).

**Extracellular thermodynamics analysis**

Calculation for ssRNA desorption from SWNT:

Johnson *et al.* (*1*) use solvent-explicit, all-atom molecular dynamics (MD) simulations for 21-nucleotide hetero- and homo-polymers adsorbing to SWNTs. In this analysis, we used the energy values for each individual nucleobase adsorption to SWNTs from Johnson *et al.* (Supplementary Table 1) to calculate the total desorption energy of each RNA sequence used in this study (Supplementary Table 2). Note that these energies are in close agreement with Das et al. (*2*), calculated from density functional theory (DFT) and experiment.

**Supplementary Table 1: Adsorption energy of each nucleotide to SWNTs (1)**

| **Base** | **Energy [kcal/mol]** |
| --- | --- |
| A | -13.84 |
| G | -14.99 |
| C | -11.07 |
| T = U | -12.68 |

Using these adsorption energies, we calculate the average desorption energy of ssRNA from the SWNT surface to be +275.3 kcal/mol ± 11.25, which we then multiply by 2 ssRNA strands to get **+550.6 kcal/mol** for both siRNA sense and antisense desorption from SWNTs (Supplementary Table 2).

**Supplementary Table 2: ssRNA-SWNT desorption energy and RNA hybridization energy for each ssRNA sequence**

| **Sequence Name** | **Sequence** | **Desorption Energy [kcal/mol]** | **Hybridization Energy**  **[kcal/mol]** |
| --- | --- | --- | --- |
| a-antisense | UUC CGU AUG UUG CAU CAC CTT | 267.1 | -39.22 |
| a-sense | GGU GAU GCA ACA UAC GGA ATT | 283.4 | -39.10 |
| b-antisense | GGG UGA AGG UGA UGC AAC ATT | 288.5 | -39.67 |
| b-sense | UGU UGC AUC ACC UUC ACC CTT | 261.5 | -39.80 |
| NT-antisense | GUA UCU CUU CAU AGC CUU ATT | 267.5 | -33.76 |
| NT-sense | UAA GGC UAU GAA GAG AUA CTT | 283.9 | -33.89 |
|  | Average | **275.3** | **-39.37** |
|  | Standard deviation | 11.25 | 0.37 |

Calculation for ssRNA to dsRNA hybridization:

ssRNA hybridization next occurs in solution, hence, we used the OligoAnalyzer tool through Integrated DNA Technologies, Inc., with the following assumed ion and RNA concentrations: [Na^+^] = 10 mM, [Mg^2+^] = 0.1mM, and [RNA] = 0.25 μM. We calculated the hybridization energy for each RNA sequence used in this study, and the average hybridization energy is **-39.37 kcal/mol** ± 0.374 (Supplementary Table 2). Therefore, the overall energy free energy change in extracellular conditions is:

$$\Delta G_{extracell}= \Delta G_{ssRNA,des}+ \Delta G_{hyb}=\left( +550.6\frac{kcal}{mol} \right)+\left( -39.373\frac{kcal}{mol} \right)=\boldsymbol{+511.2}\frac{\boldsymbol{kcal}}{\boldsymbol{mol}}$$

Based on this positive, unfavorable free energy change, our analysis demonstrates that it is unlikely for RNA desorption and hybridization to take place spontaneously in the extracellular environment when both complementary RNA strands are initially adsorbed on SWNTs.

**Intracellular thermodynamics analysis**

The intracellular environment is crowded with biomolecules, is highly dynamic, and intracellular components of the cell cytoplasm such as proteins and lipids are known to bind to SWNTs (*3*). Accordingly, once inside the cell, cytoplasmic biomolecules will likely to adsorb on SWNTs, as observed by SWNT solvatochromic shifts (**Fig. S3**), thus favoring RNA desorption in the presence of a complementary polynucleotide sequence (*4*). Therefore, we hypothesized that inside the cell, RNA desorption and hybridization are likely to take place, lowering the free energy of “bare” SWNTs by direct biomolecule replacement, hence making this process thermodynamically favorable under cytoplasmic conditions as calculated below (**Fig. 3**). We assume that the same ssRNA desorption energies apply, but now this unfavorable ssRNA desorption is countered by favorable protein adsorption energy to the SWNT surface.

Calculation for protein adsorption energy to SWNTs:

We assume that the RNA desorption and protein adsorption steps are taking place independently. Shen *et al.* (*5*) use MD simulation for the adsorption of human serum albumin (HSA) helices on different chirality SWNTs with water solvent, and calculate an average protein adsorption energy of -14.51 kcal/mol ± 1.858 per residue. Similarly, DFT calculations have reported protein adsorption to carbon nanotubes with an average energy of ~ **-10 kcal/mol per residue** (*6*, *7*). The median number of residues per protein in *A. thaliana* is 356 residues (*8*). We assumed -10 kcal/mol binding energy per residue and that only 10% of the residues participate in the adsorption process to the SWNT surface. Accordingly, we obtained -356.0 kcal/mol for an average protein adsorption energy to SWNTs. Note that this energy is within a reasonable order of magnitude, as HSA adsorption on carboxylated SWNTs is ~ -500 kcal/mol via MD simulation (*9*). Then by multiplying by 2 SWNTs, we get **-712.0 kcal/mol** for protein adsorption to the SWNT surface. Therefore, the overall change in free energy in intracellular conditions is:

$$\Delta G_{intracell}= \Delta G_{ssRNA,des}+ \Delta G_{prot,ads}+\Delta G_{hyb}$$

$$=\left( +550.6\frac{kcal}{mol} \right)+\left( -712.0\frac{kcal}{mol} \right)+\left( -39.373\frac{kcal}{mol} \right)\boldsymbol{=-200.8}\frac{\boldsymbol{kcal}}{\boldsymbol{mol}}$$

Our analysis shows that this overall free energy change is negative inside plant cells, which demonstrates that RNA desorption from the SWNT surface and subsequent hybridization of complementary siRNA sequences are favorable and spontaneous in intracellular conditions, recapitulating our experimental results.

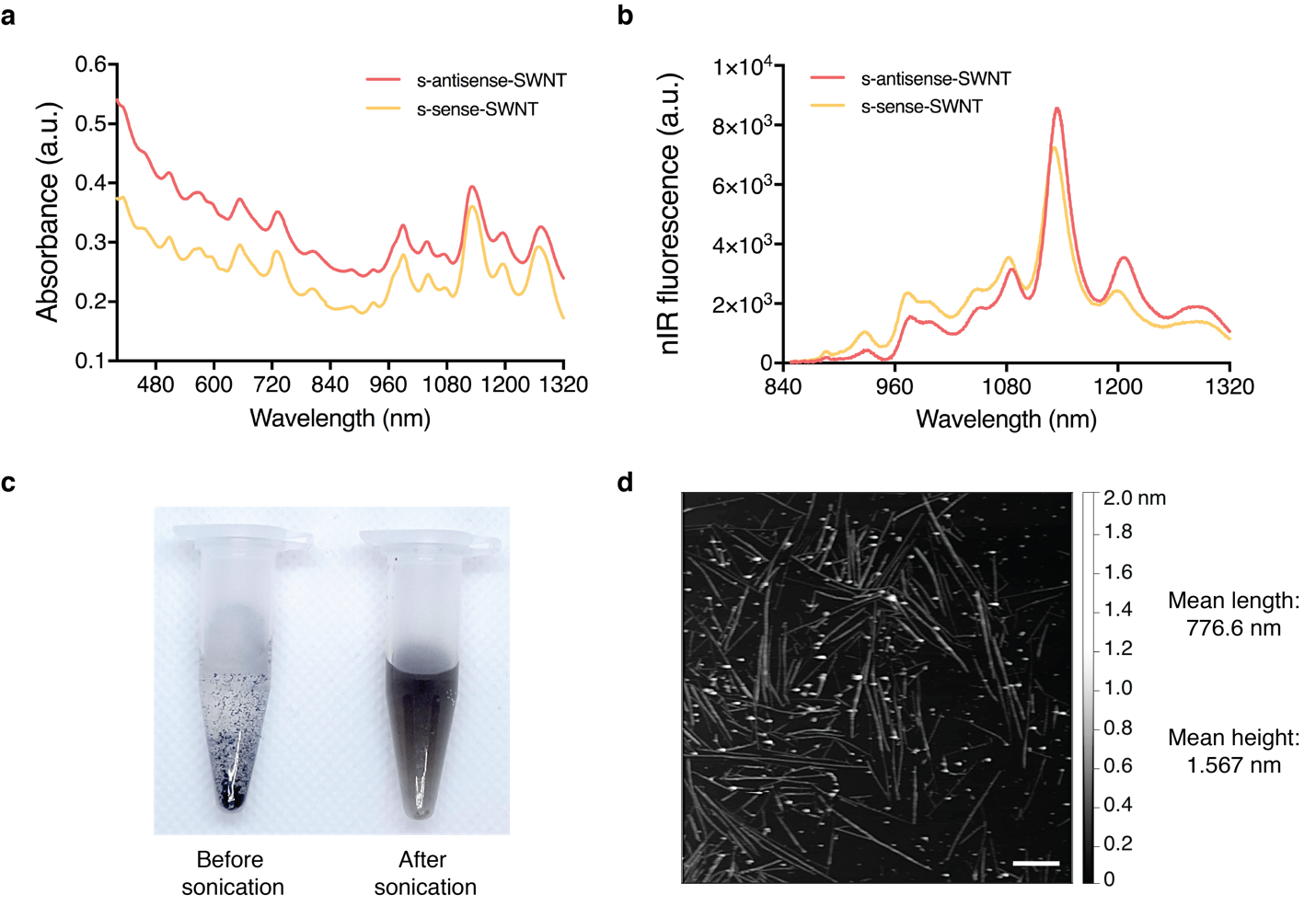

**Supplementary Fig. 1. Non-targeting s-RNA-SWNT suspension characterization and AFM imaging. a** Absorbance spectra of s-antisense-SWNT and s-sense-SWNT suspensions. **b** Near-infrared (nIR) fluorescence spectra of s-antisense-SWNT and s-sense-SWNT suspensions. **c** Photo on the left showing RNA SWNT mixture before probe-tip sonication where unsuspended SWNTs aggregate due to van der Waals and hydrophobic interactions between SWNTs, and photo on the right showing homogenous dark-colored individually suspended RNA-SWNT solution that is colloidally stable following probe-tip sonication. **d** Representative atomic force microscopy (AFM) image of an ssRNA-SWNT suspension. Scale bar, 100 nm. Mean length of RNA-SWNTs is 776.6 nm (st. dev. 163 nm) and mean height is 1.567 nm (st. dev. 0.38 nm) for N = 25 SWNTs.

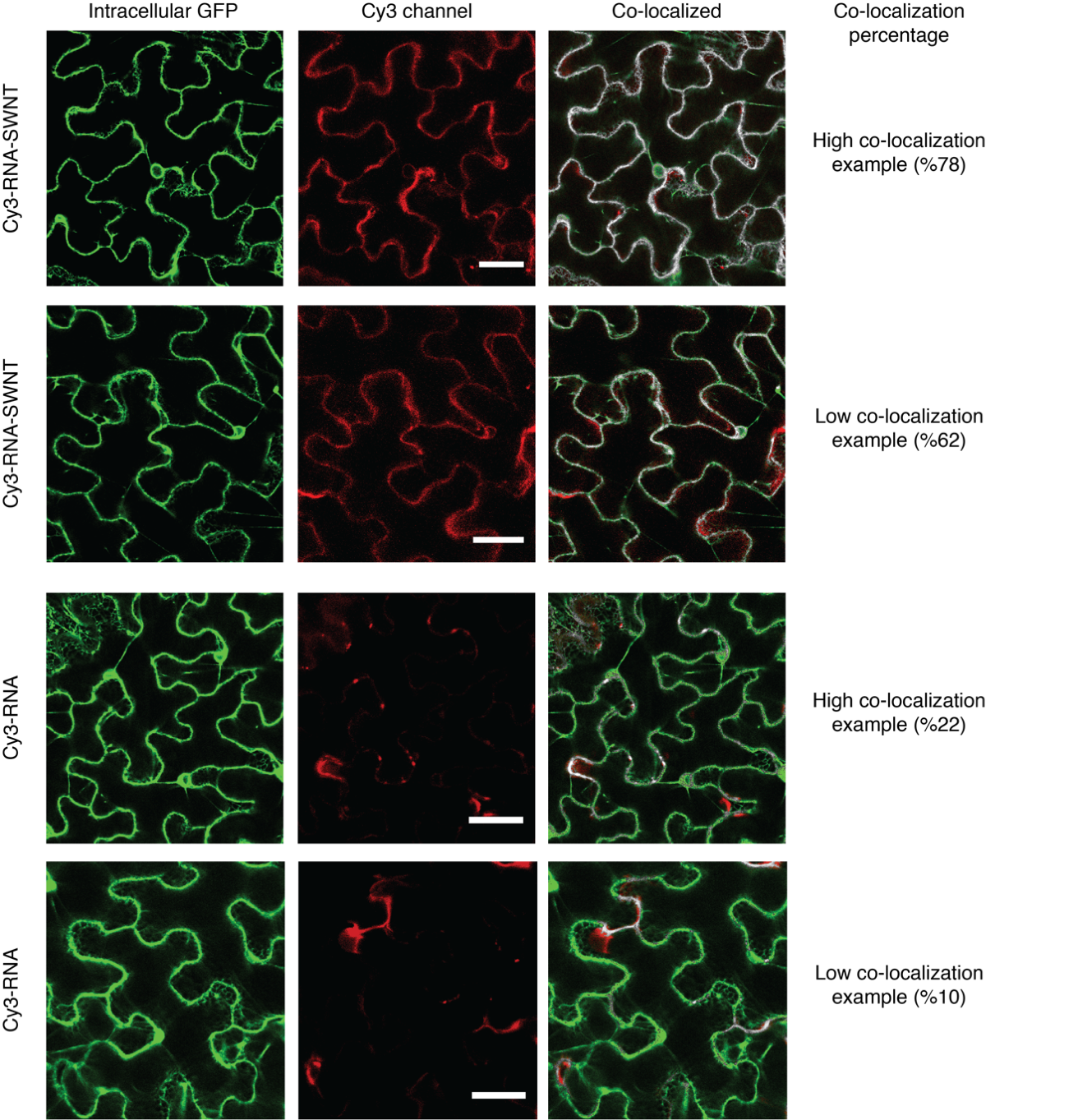

**Supplementary Fig. 2. Internalization of Cy3-RNA-SWNT and Cy3-RNA into mGFP5 *Nicotiana benthamiana* leaves assessed with confocal microscopy and co-localization analysis.** Top two rows: Cy3-RNA-SWNT infiltrated *Nb* leaf images showing examples of high (78%) and low (62%) co-localization percentages of intracellular GFP with Cy3-RNA-SWNT fluorescence. Bottom two rows: Free Cy3-RNA infiltrated *Nb* leaf images showing examples of high (22%) and low (10%) co-localization percentage of intracellular GFP with free Cy3-RNA fluorescence. Scale bars are 40 µm.

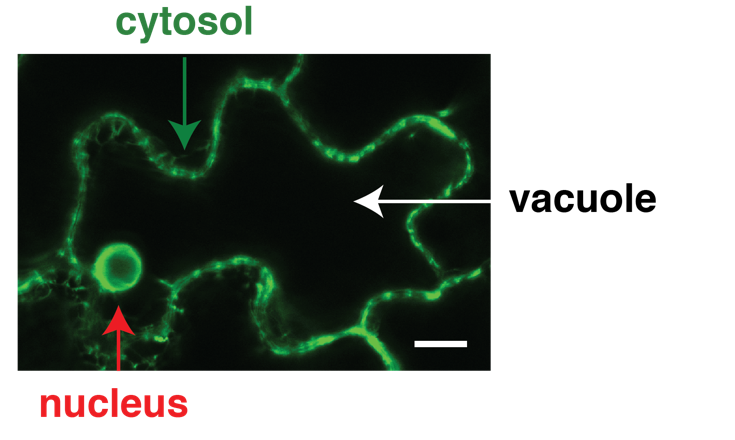

**Supplementary Fig. 3. Subcellular areas in GFP *Nicotiana benthamiana* leaf cells.** Green arrow shows the cytosol pushed back to the cell membrane due to the presence of the large central plant cell vacuole. Red arrow marks the nucleus and white arrow marks the vacuole in a leaf epidermal cell imaged with confocal microscopy. Scale bar: 20 µm.

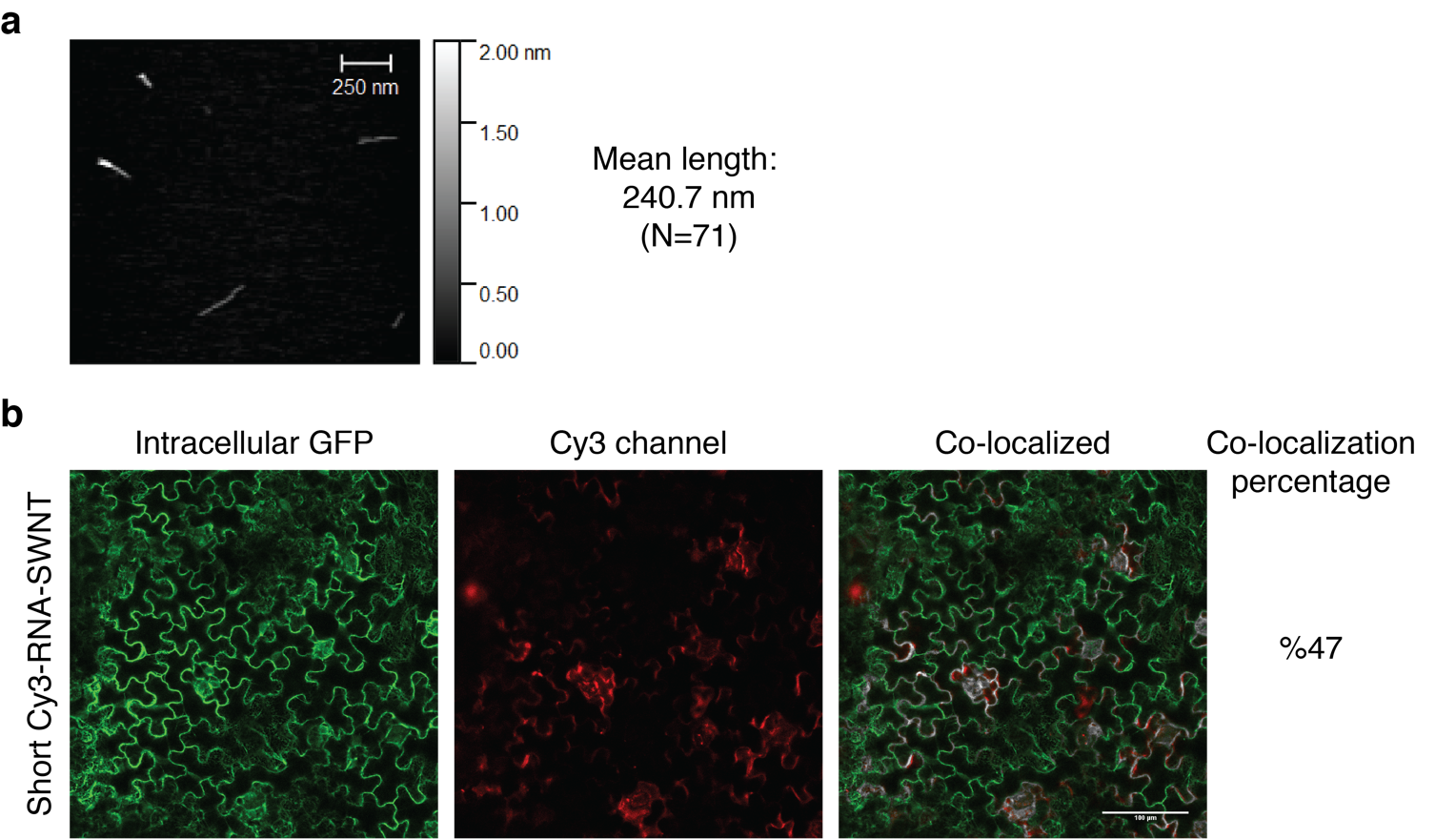

**Supplementary Fig. 4. Short Cy3-RNA-SWNTs and their internalization efficiency analysis. a** Representative AFM image showing shortened SWNTs with an average length of ~250 nm. **b** Representative short Cy3-RNA-SWNT infiltrated *Nb* leaf confocal image showing 47% co-localization percentage of intracellular GFP with Cy3-RNA-SWNT fluorescence.

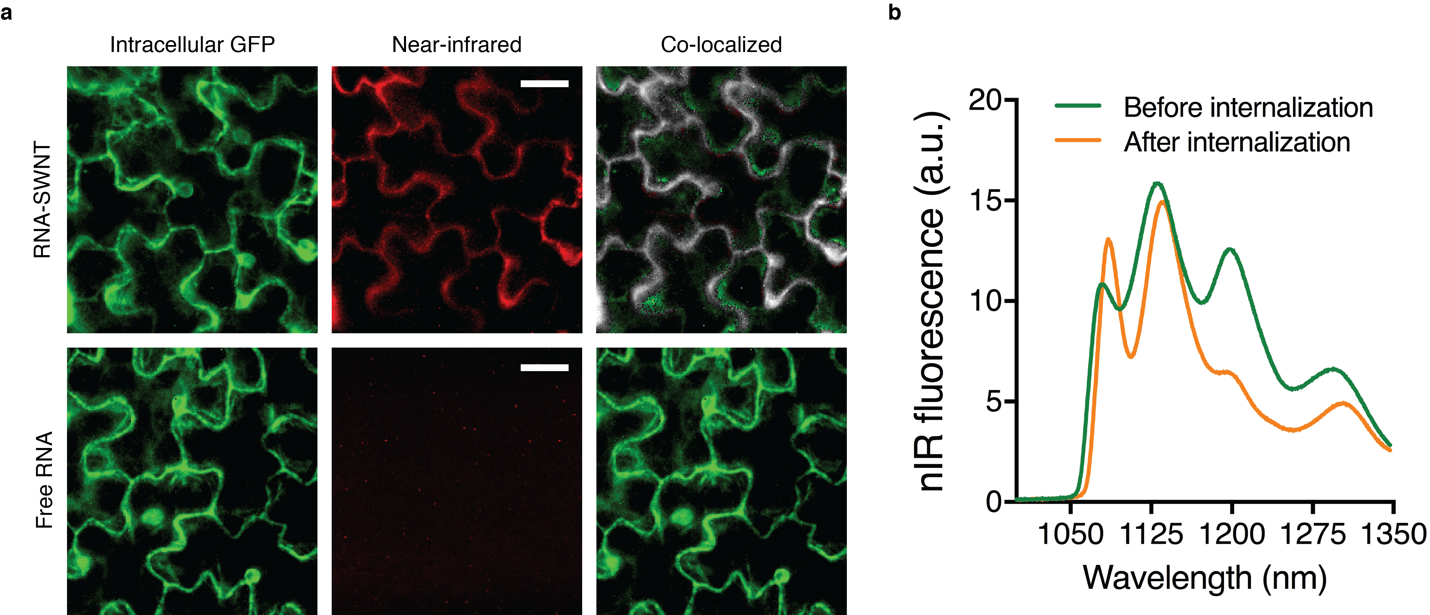

**Supplementary Fig. 5. nIR imaging shows internalization of RNA-SWNT suspensions into mGFP5 *Nicotiana benthamiana* leaves. a** Top row: RNA-SWNT infiltrated *Nb* leaf images showing high co-localization efficiency of intracellular GFP with intrinsic nIR SWNT fluorescence. Bottom row: Free RNA infiltrated *Nb* leaf images showing no co-localization of free RNA inside cells. Scale bars are 20 µm. **b** nIR fluorescence spectra of RNA-SWNTs before and after internalization into leaf cells. Spectra were obtained with a 1050-nm long pass filter to avoid the autofluorescence of chlorophyll from leaves.

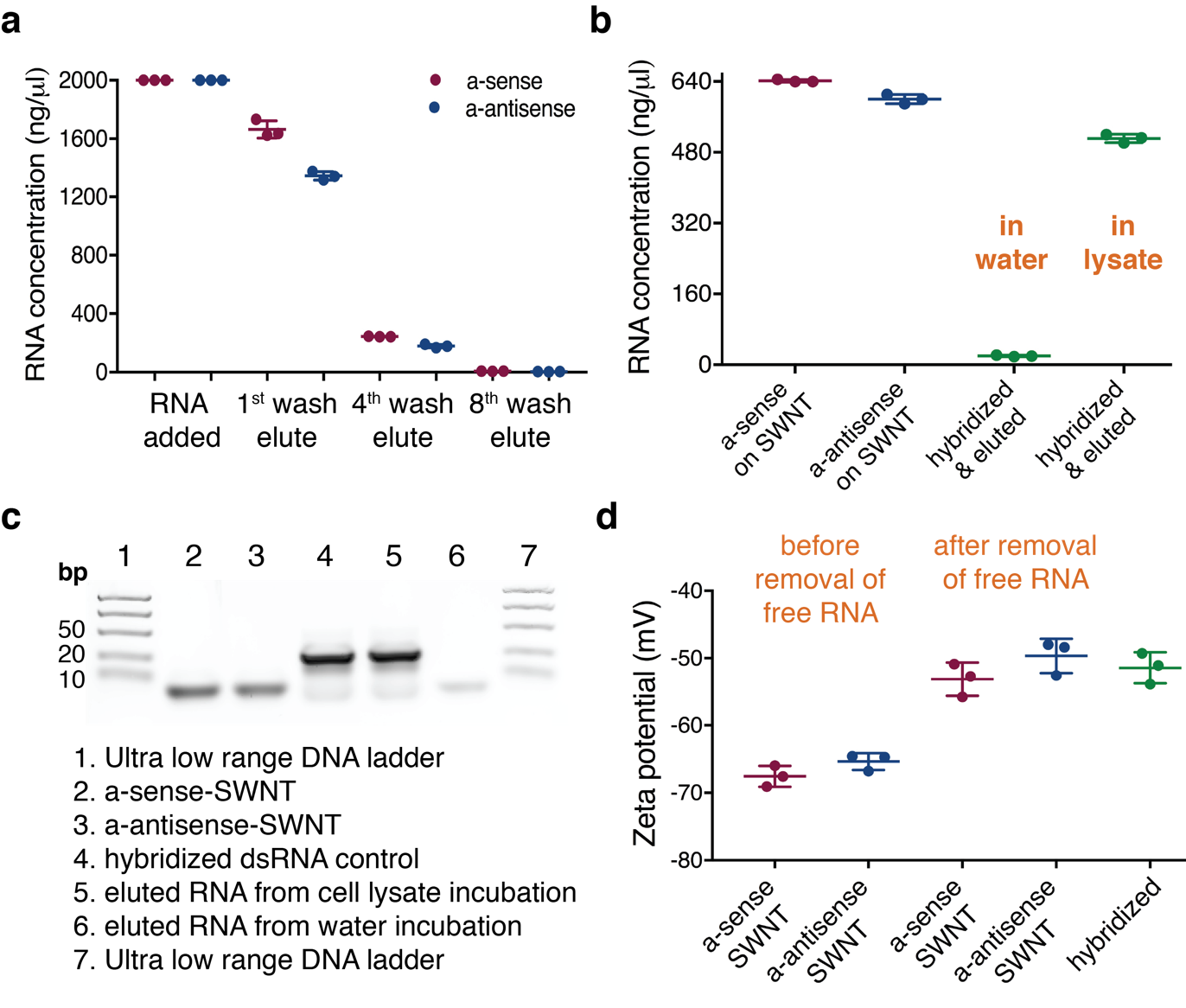

**Supplementary Fig. 6. Characterization of dsRNA hybridization and desorption. a** a-sense and a-antisense RNA amount added to suspend SWNTs, RNA concentration in the 1^st^, 4^th^ and 8^th^ flow-through solutions after centrifugation with 100K spin filters to remove free RNA after suspension with SWNTs (RNA concentration is calculated via absorbance measurements at 260 nm). **b** RNA amount on suspended SWNTs (calculated via total RNA added – free RNA removed via spin filtration), dsRNA eluted after hybridization in water and in cell lysate conditions. **c** 4% agarose gel showing eluted RNA from cell lysate and water incubations, 97% double stranded RNA is eluted from the cell lysate incubation and 19% of single stranded RNA is eluted from the water incubation sample. **d** Zeta potential measurements for a-sense-SWNT and a-antisense-SWNT suspensions before and after the removal of free RNA, and after mixing a-sense-SWNT and a-antisense-SWNT solutions in water and eluting free RNA (hybridized).

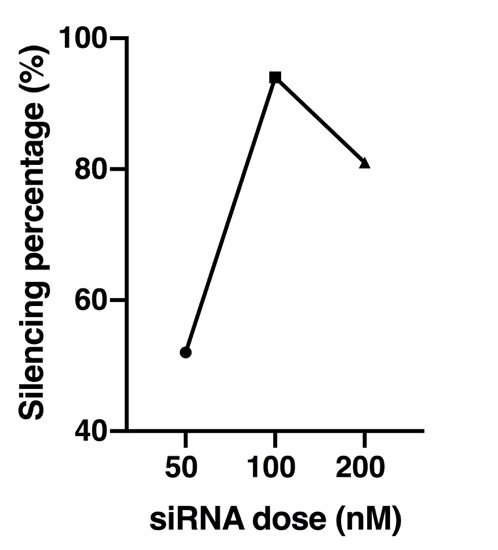

**Supplementary Fig. 7. Optimization of siRNA dose on SWNTs for *mGFP5* silencing.** Final siRNA concentration of 50, 100 and 200 nM on SWNTs, and corresponding gene silencing efficiencies at 1-day post-infiltration measured via qPCR assay.

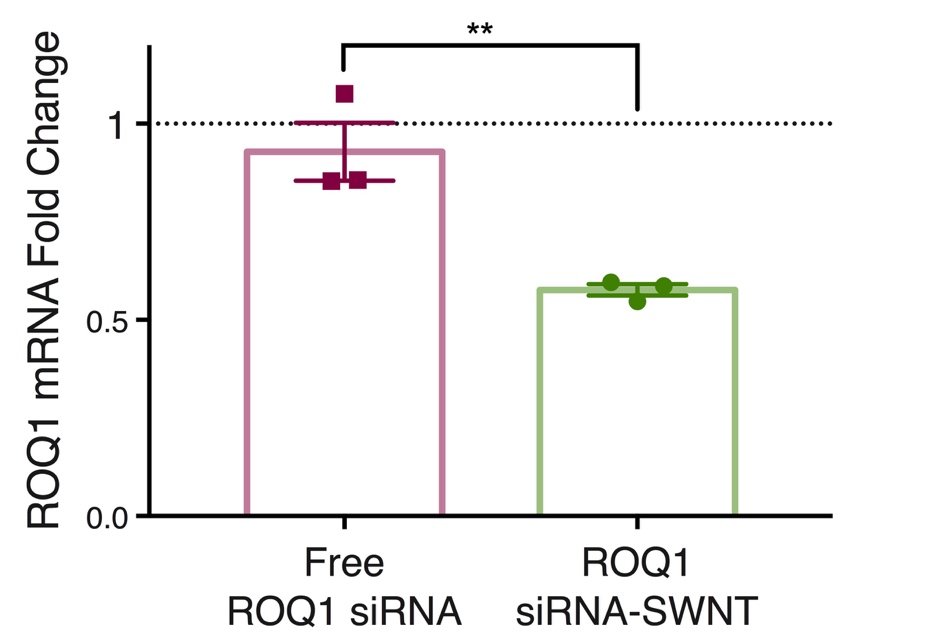

**Supplementary Fig. 8. Silencing of endogenous functional *Nicotiana benthamiana* ROQ1 gene with siRNA-SWNTs.** Free ROQ1 siRNA without SWNTs do not show significant silencing of ROQ1 gene, whereas 100 nM ROQ1 siRNA on SWNTs yields nearly 50% mRNA reduction at Day 1 as assessed by qPCR of infiltrated *Nicotiana benthamiana* leaves compared to the non-treated control leaves. ** P = 0.0094 in two-tailed t-test. Error bars indicate s.e.m. (n = 3).

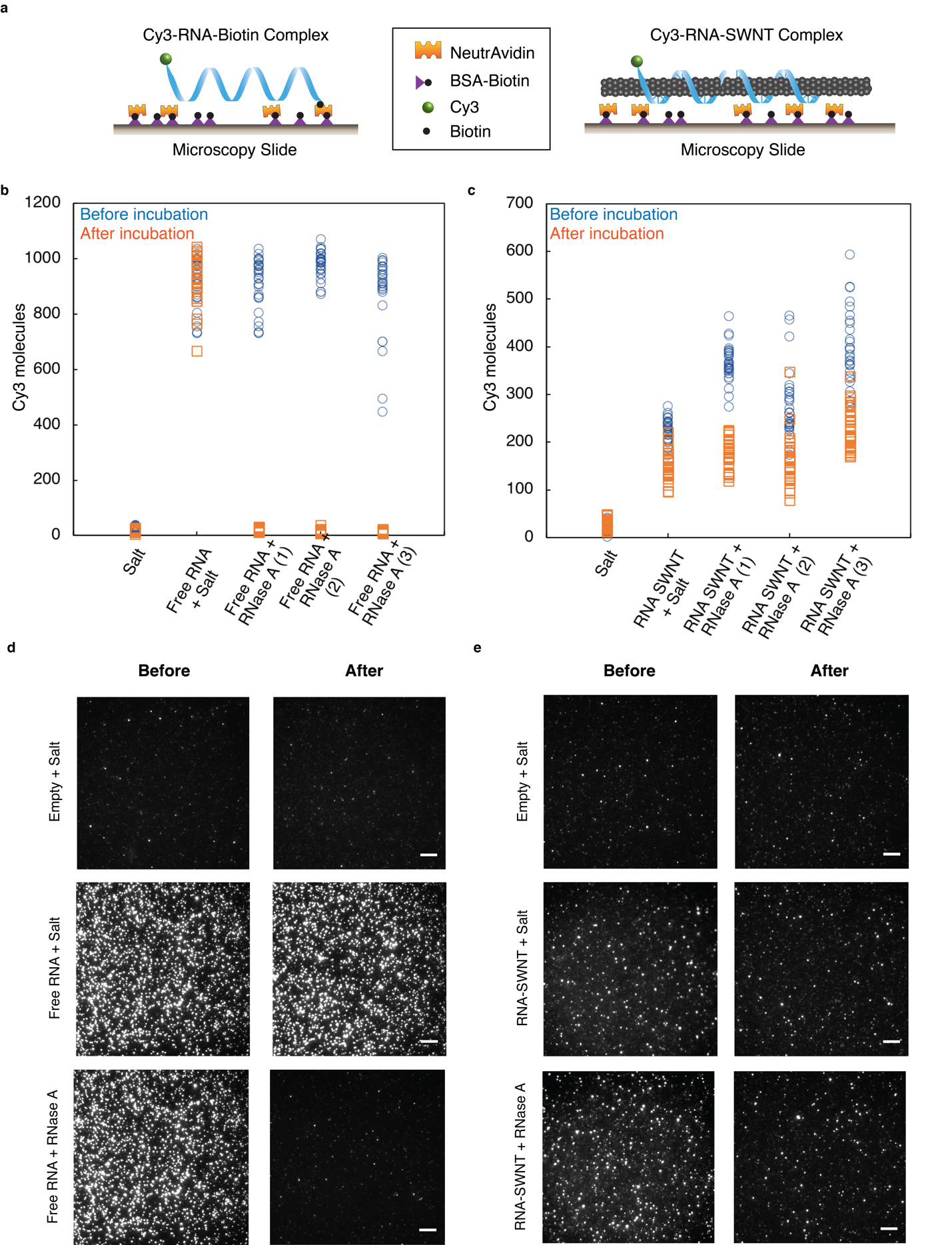

**Supplementary Fig. 9. Single molecule TIRF (smTIRF) microscopy demonstrates RNA protection from nuclease degradation when adsorbed to SWNTs. a** Schematics of microfluidic slides for immobilization and smTIRF imaging of Cy3-RNA-Biotin and Cy3-RNA-SWNT complexes. **b** Raw smTIRF data for empty channel rinsed with salt solution, free RNA incubated with salt solution, and three experimental replicates of free RNA incubated with RNase A, blue: before incubation and orange: after incubation. Data from 30 fields of views was plotted for each sample before and after treatment. **c** Raw smTIRF data for empty channel rinsed with salt solution, RNA-SWNT incubated with salt solution, and three experimental replicates of RNA-SWNT incubated with RNase A, blue: before incubation and orange: after incubation with RNase A. Data from 30 fields of views was plotted for each sample before and after treatment. **d** Representative TIRF microscopy images for each sample of free RNA before and after incubation with salt and RNase A. **e** Representative TIRF microscopy images for each sample of RNA-SWNTs before and after incubation with salt and RNase A. All scale bars, 5 µm.

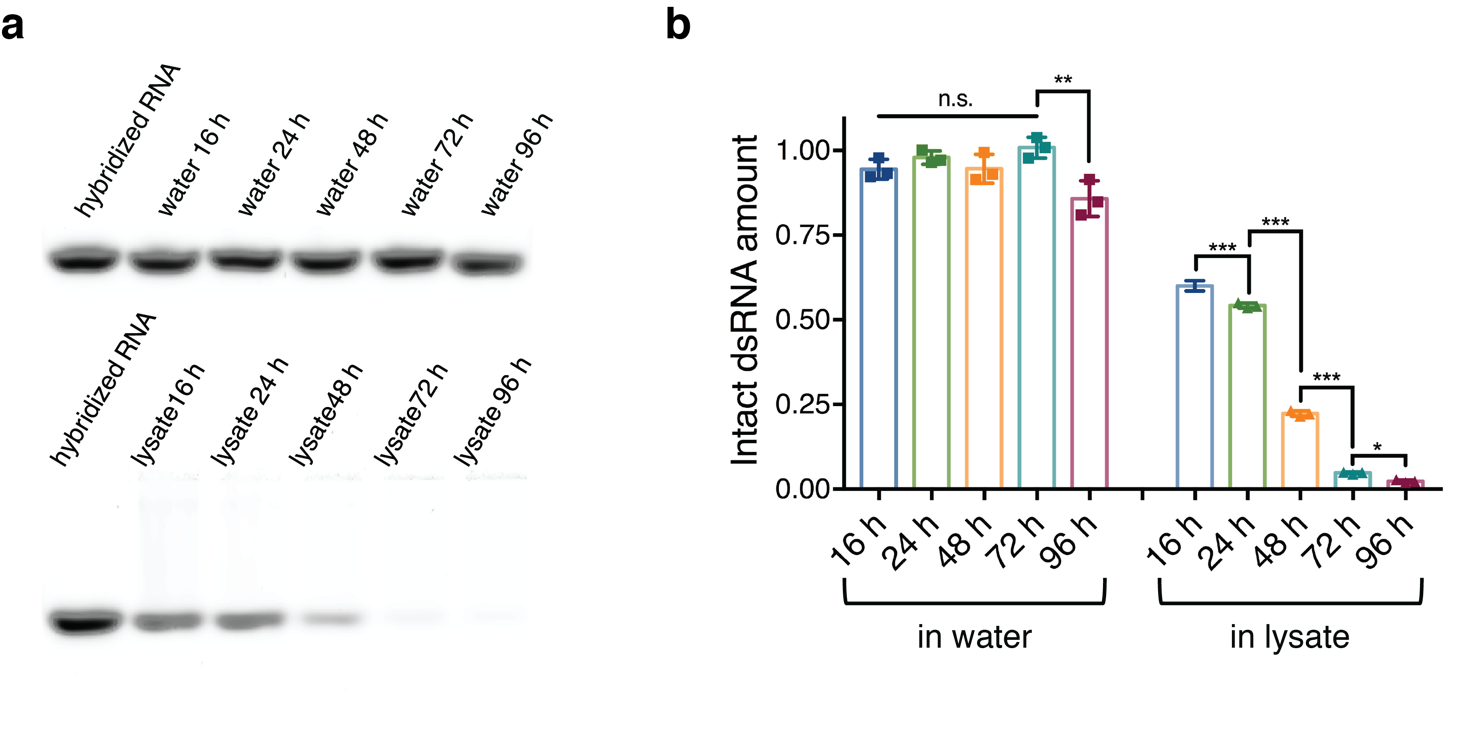

**Supplementary Fig. 10. dsRNA stability in cell lysate. a** Hybridized double stranded siRNA samples incubated in nuclease-free water and cell lysate solutions at room temperature for 16, 24, 48, 72 and 96 hours, and run on a 2% agarose gel. **b** Quantification of RNA bands from the gel in part a and two other replicates, using the Image J gel analyzer tool. All band intensities are normalized with respect to the hybridized RNA band intensity at time zero without any treatment.

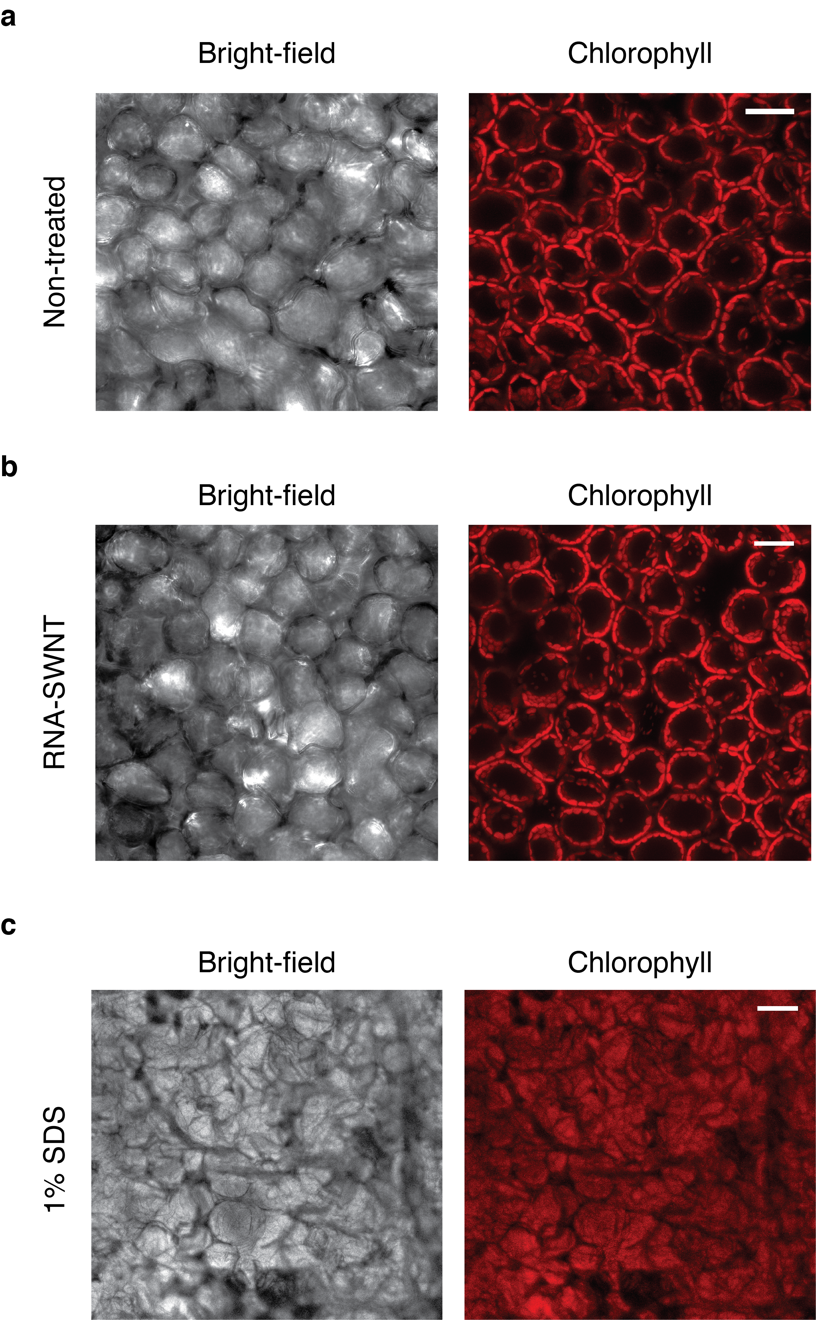

**Supplementary Fig. 11. Confocal microscopy imaging of the *Nb* leaf tissue to assess cellular damage.** **a** Representative bright-field and chlorophyll images of non-treated *Nb* leaf. **b** RNA-SWNT treated *Nb* leaf and **c** 1% SDS treated *Nb* leaf as a positive control of leaf tissue damage. All scale bars, 20 µm.

**Supplementary Table 3: RNA sequences and primers used in this study**

| **RNA sequences:** | (all sequences written as 5’ to 3’) |
| --- | --- |
| a-antisense | UUC CGU AUG UUG CAU CAC CTT |
| a-sense | GGU GAU GCA ACA UAC GGA ATT |
| b-antisense | GGG UGA AGG UGA UGC AAC ATT |
| b-sense | UGU UGC AUC ACC UUC ACC CTT |
| s-antisense | GUA UCU CUU CAU AGC CUU ATT |
| s-sense | UAA GGC UAU GAA GAG AUA CTT |
| Cy3 tagged a-antisense | Cy3/UUC CGU AUG UUG CAU CAC CTT |
| ROQ1-sense | GGU UUA AUU UGG UGU AUA A |
| ROQ1-antisense | UUA UAC ACC AAA UUA AAC C |
| **Primers for qPCR:** |  |
| EF1 forward | TGG TGT CCT CAA GCC TGG TAT GGT TG |
| EF1 reverse | ACG CTT GAG ATC CTT AAC CGC AAC ATT CTT |
| mGFP5 forward | AGT GGA GAG GGT GAA GGT GAT G |
| mGFP5 reverse | GCA TTG AAC ACC ATA AGA GAA AGT AGT G |
| NbrbohB forward | TTT CTC TGA GGT TTG CCA GCC ACC ACC TAA |
| NbrbohB reverse | GCC TTC ATG TTG TTG ACA ATG TCT TTA ACA |
| ROQ1 forward | TCC CCG ACA TAA AGG AAT GC |
| ROQ1 reverse | GTC CCC TGG ACT CAA ACA GG |
